# Innate liking and disgust reactions elicited by intraoral capsaicin in male mice

**DOI:** 10.1101/2024.04.21.590483

**Authors:** Yibin Han, Daisuke H. Tanaka, Naofumi Uesaka

## Abstract

Liking and disgust are the primary positive and negative emotions, respectively, and are crucial for nutrient intake and toxin avoidance. These emotions are induced by multimodal stimuli such as taste, olfactory, and somatosensory inputs, and their dysregulation is evident in various psychiatric disorders. To understand the biological basis of liking and disgust, it is crucial to establish an animal model that allows for quantitative estimation of liking and disgust in response to multimodal stimuli. The only readout shared by many species, including humans, for liking and disgust, has been taste reactivity. However, readouts of non-taste stimuli-induced emotions remain unestablished. Here, we show that intraoral administration of capsaicin, a chemosomatosensory stimulus, elicits orofacial and bodily reactions in male mice similar to those observed in taste reactivity. Capsaicin induced liking reactions at low concentrations and disgust reactions at high concentrations. Capsaicin-induced disgust reactions consisted of various reactions, including gape and forelimb flail, with the proportion of each reaction among the disgust reactions being similar to that induced by bitter and sour stimuli. These findings indicate that orofacial and bodily reactions, defined as taste reactivity, are elicited not only by taste stimuli, but also by intraoral chemosomatosensory stimuli. Understanding the biological basis of capsaicin-induced orofacial and bodily reactions will advance our understanding of the fundamental mechanisms underlying liking and disgust across sensory modalities.

## Introduction

The emotional responses of liking and disgust are fundamental to many animals, including humans, and influence our daily lives and well-being. Liking, or pleasure, is one of the primitive positive emotions elicited by the taste of sweet food or affectionate contact, and is thought to have evolved to promote nutrient intake and reproduction (Berridge and Kringelbach, 2008; 2015). Disgust is one of the basic negative emotions induced by the distaste of certain foods and contact with feces and is believed to have evolved as a protective mechanism against toxins and pathogens (Darwin, 1872/1965; Rozin and Fallon, 1987; Curtis et al., 2004; Chapman and Anderson, 2012; Tybur et al., 2013).

In humans, liking and disgust are induced by external stimuli of all five modalities: taste (Steiner, 1979), olfactory (Engen and McBurney, 1964), somatosensory (Rozin and Schiller, 1980), auditory (Scott et al., 1997), and visual stimuli (Adolphs et al., 1994). The expression of liking and disgust is dysregulated in various psychiatric disorders, such as anhedonia, in which liking is diminished, and is associated with depression (Gorwood, 2008), Parkinson’s disease (Loas et al., 2012), and schizophrenia (Der-Avakian and Markou, 2012; Lambert, 2018), while excessive disgust is associated with depression, anxiety, and obsessive-compulsive disorder (Davey, 2011).

To understand the biological basis of liking and disgust in health and disease, it is crucial to establish a quantitative method to estimate the degree of liking and disgust in animals. The taste reactivity test is the gold standard method for quantitatively estimating the degree of liking and disgust in animals and quantifies orofacial and bodily reactions consisting of orofacial, head, and upper-limb motor reactions induced by taste stimuli (Grill and Norgren, 1978; Berridge, 2000). Orofacial and bodily reactions defined in the taste reactivity test are highly conserved in mammals, including humans, monkeys, and rodents, and are widely used as reliable indicators of liking and disgust induced by taste stimuli (Berridge, 2000). However, the stimuli used to date in this test have been limited to taste stimuli, and no method has been established to estimate the liking or disgust induced by the stimulation of other sensory modalities.

The trigeminal sensory nerve contains C-and A-delta-fibers that convey nociceptive signals from the oral cavity (Edvisson et al., 2019). C-and A-delta-fibers are excited by capsaicin, a ligand for the transient receptor potential vanilloid 1 receptor (LaMotte et al., 1992; Caterina et al., 2000; Ringkamp et al., 2001). Capsaicin is widely used as a noxious somatosensory stimulus in pain research (Winter et al., 1995; Ngom et al., 2001; Baad-Hansen et al., 2003; O’Neill et al., 2012; Zhao et al., 2012; Lu et al., 2013; Naganawa et al., 2015). In humans, intraoral administration of capsaicin can elicit both liking and disgust at varying concentrations, with pain induced by high concentrations, although there are significant individual differences (Rozin and Schiller, 1980; Rozin et al., 1982; Stevenson and Yeomans, 1993; 1995; Tornwall et al., 2012; Byrnes and Hayes, 2016). In rodents, high concentrations of capsaicin are avoided, with a preference for water over any capsaicin concentration (Bachmanov et al., 1996; Simons et al., 2001; Furuse et al., 2002; Narukawa and Misaka, 2021). However, it remains unclear whether intraoral capsaicin induces liking and disgust reactions, which are defined as taste reactivity.

In this study, we examined orofacial and bodily reactions representing liking and disgust using three different concentrations of intraoral capsaicin in male mice and compared them with those induced by two representative tastants, bitter quinine and sour citric acid.

## Materials and methods

### Animals

Adult (6-7 weeks-old) male C57BL/6J mice (*n*=16) were obtained from Japan SLC Inc. (Shizuoka, Japan) and used in the present study. Upon arrival, mice were housed in clear plastic cages (18 × 26 × 13 cm) with cellulose fiber chips (Alpha-Dri; Shepherd Specialty Papers, Kalamazoo, MI, USA) in groups of 2-4 males per cage and were then housed individually in smaller cages (14 × 21 × 12 cm) immediately before handling. Mice were maintained at 22 ± 1 °C under a 12 h light/dark cycles (lights on at 8:00 am) and given *ad libitum* access to food (CE-2; CLEA Japan Inc., Tokyo, Japan) and tap water. The tap water was changed to Milli-Q water 3 days before beginning the reaction experiments. All reaction experiments were performed in the light cycle. All the animal experiments were approved (No. A2022-005C6) by the Institutional Animal Care and Use Committee of Tokyo Medical and Dental University, and performed in accordance with the relevant guidelines and regulations.

### Surgery

Intraoral tube implantation was performed as previously described (Tanaka et al., 2019), with minor modifications. Mice were anesthetized by intraperitoneal injection of a mixture of midazolam (5.2 mg/kg body weight (BW), Dormicum; Astellas Pharma Inc., Tokyo, Japan), butorphanol (6.5 mg/kg BW, Vetorphale; Meiji Animal Health Co., Ltd., Kumamoto, Japan), and medetomidine (0.39 mg/kg BW, Domitor; Nippon Zenyaku Kogyo Co., Ltd., Fukushima, Japan). The depth of anesthesia was maintained at a sufficient level to prevent paw pinch reflex. The eyes were protected using ophthalmic ointment (Tarivid; Santen, Osaka, Japan). The epidermis of the parietal was cut into elliptical shapes and removed. A curved needle attached to an intraoral polyethylene tube (SP-10; Natsume Seisakusho Co., Ltd., Tokyo, Japan) was inserted from the incision site and advanced subcutaneously posterior to the eye, to exit at a point lateral to the first maxillary molar on the right side of the mouth. The intraoral end of the tube was heat-flared to an approximate diameter of 1 mm to prevent it from drawing into the oral mucosa. Mice were mounted using ear bars in a stereotaxic frame (Stoelting Co., Wood Dale, IL, USA). A piece of plastic bar was fixed to the skull using dental cement (Super-Bond C&B; Sun Medical, Shiga, Japan). The mice received subcutaneous injections of the antibiotic chloramphenicol sodium succinate (60 mg/kg BW, Chloromycetin Succinate; Daiichi Sankyo Co., Ltd., Tokyo, Japan) for infection prevention and carprofen (5 mg/kg BW, Rimadyl; Zoetis, Tokyo, Japan) for pain relief, and received intraperitoneal injection of atipamezole (0.39 mg/kg BW, Antisedan; Noppon Zenyaku Kogyo Co., Ltd.) for reversing the anesthetic effects of medetomidine.

Three days after surgery, one end of a delivery tube (SP-10; Natsume Seisakusho Co., Ltd.) was connected to the end of the intraoral tube fixed to the plastic bar on the mouse head using a connector (KN-394, two directions (0.3+0.3); Natsume Seisakusho Co., Ltd.), and the other end was connected to a needle (30Gx13mm; ReactSystem Co., Osaka, Japan) attached to a 1-mL syringe, and ∼20 µL of Milli-Q water was infused into the mouth of the animal to test the patency of the tubes. The intraoral tube was flushed with approximately 20 µL Milli-Q water every weekday to prevent occlusion. The mice were allowed 1-2 weeks to recover from surgery before beginning the reaction experiments.

### Preparation of stimulant solutions

Capsaicin (code#07127-21; nacalai tesque, Kyoto, Japan) was dissolved in ethanol to prepare a 660 mM capsaicin solution, then it was diluted with Milli-Q water to make a 330 µM capsaicin solution containing 0.05% ethanol. The 330 µM capsaicin solution was further diluted with Milli-Q water to obtain 33 µM and 3.3 µM capsaicin solutions. The final concentration of ethanol in all three capsaicin solutions was adjusted to 0.05% by adding ethanol appropriately. Tannic acid (code#32616-02; nacalai tesque) was dissolved in Milli-Q water to prepare a 75 mg/mL solution, which was then diluted with Milli-Q water to prepare 7.5 mg/mL and 0.75 mg/mL solutions. Citric acid (code#09109-85; nacalai tesque) was dissolved in Milli-Q water to prepare a 1 M solution, and diluted with Milli-Q water to make 0.1 M and 0.01 M solutions. The pH values for the 1 M, 0.1 M, and 0.01 M citric acid solutions were approximately 1.5, 2, and 2.5, respectively. Quinine (code#22630; Sigma-Aldrich Co., MO, USA) was dissolved in Milli-Q water to prepare a 10 mM solution, which was then diluted with Milli-Q water to obtain 1 mM and 0.1 mM solutions.

To investigate the differences in the disgust reactions elicited by various stimulus solutions, we prepared three concentrations of each solution, anticipating a gradual change in the reaction magnitude. Initially, based on previous studies (Galaverna et al., 1993; Bachmanov et al., 1996; Simons et al., 2001; Furuse et al., 2002; Tanaka et al., 2019; 2021), we conducted preliminary experiments to determine the concentration at which disgust reactions begin to be induced. However, at concentrations of 33 µM, 1 mM, and 0.1 M, respectively, consistent induction of disgust was noted. Subsequently, to plot the results on a graph with the logarithm of the stimulus solution concentration on the x-axis and the frequency of disgust reactions on the y-axis, we prepared three concentrations for each solution: the concentration at which disgust reactions began to be observed, one-tenth of that concentration, and ten times that concentration. Consequently, in this study, we used solutions of 3.3, 33, and 330 µM capsaicin; 0.1, 1, and 10 mM quinine; and 0.01, 0.1, and 1 M citric acid. Indeed, the disgust reactions at the highest concentration of each solution were significantly greater than those at the lower concentrations (Fig. 2), which we believe was necessary to clearly capture the disgust reactions induced by each solution. However, it is possible that concentrations higher than the medium, but lower than the high concentration used in this study, could also induce clear disgust reactions. In addition, since it has been reported that a 0.3M citric acid stimulus elicits responses in lingual trigeminal fibers (Bryant and Moore, 1995), it is possible that the 1M citric acid stimulus used in the present study may be transmitted not only as a taste signal but also as a chemosomatosensory signal.

### Taste reactivity tests for comparison among capsaicin, quinine, and citric acid

For these tests (all figures except for Fig. 1D-G and Supplemental fig.1), eight mice underwent oral surgery. All eight mice were used for the capsaicin test, and six of the eight mice were used for the quinine and citric acid tests. We used one–two mice in each independent experiment and conducted six independent experiments. Definition of "independent experiment": The term "independent experiment" refers to a complete experimental series starting from surgery through all related tests. For instance, if two mice undergo surgery and complete all tests in one temporal block, this would be counted as "n=2 mice from 1 independent experiment." In the first two independent experiments, we used two mice in total and focused on their responses to capsaicin; therefore, experiments with quinine and citric acid were not conducted. In the remaining four independent experiments, six mice were used in total, and the experiments were conducted with quinine and citric acid in addition to capsaicin. Therefore, data on responses to capsaicin stimuli were obtained from eight mice across six independent experiments. Data for quinine and citric acid stimuli were obtained from six mice across four independent experiments.

**Figure 1.**
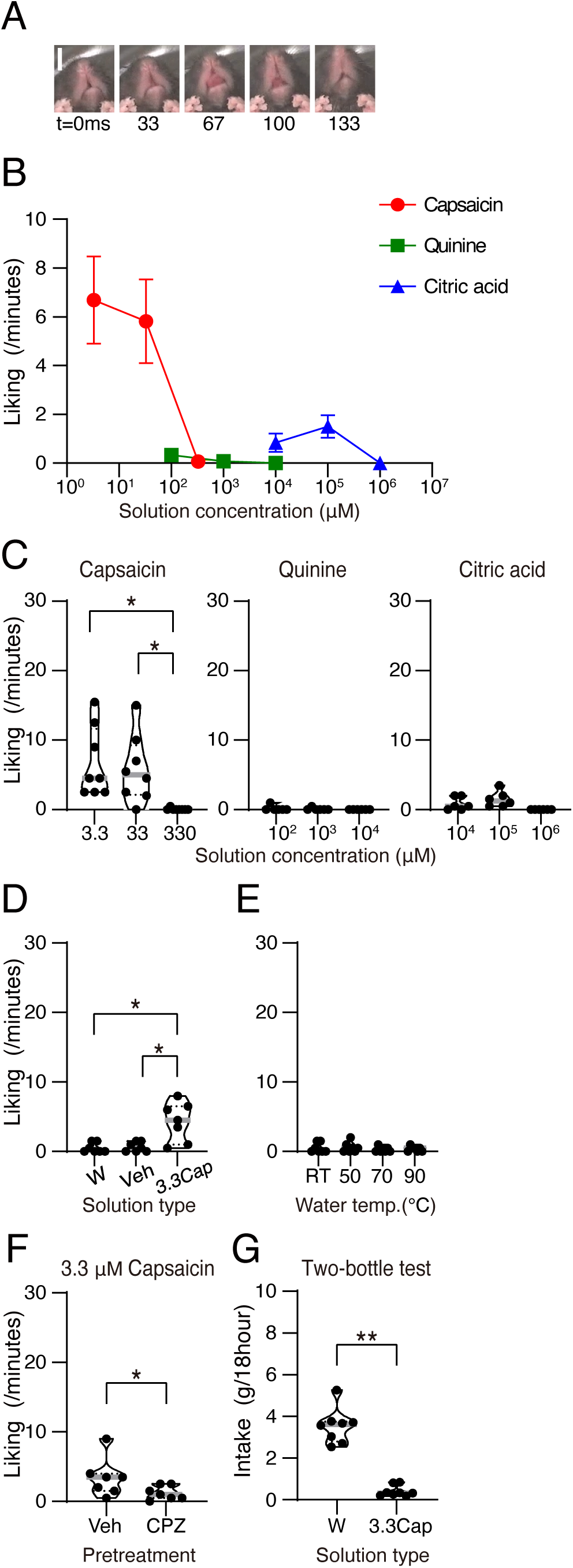
Capsaicin of low and medium concentrations elicits liking reactions. **(A)** Representative time-lapse sequences of tongue protrusion during the intraoral infusion of capsaicin. The numbers at the bottom of each panel indicate the elapsed time (milliseconds). Scale bar, 5 mm. **(B)** Relationship between liking reaction scores during intraoral infusion of capsaicin (red circles; *n* = 8 mice from six independent experiments), quinine (green rectangles; *n* = 6 mice from four independent experiments), and citric acid (blue triangles; *n* = 6 mice from four independent experiments), and their concentrations. Error bars are SEM. **(C)** The score for liking reactions during intraoral infusion of three different concentrations of capsaicin (left panel,l; *n* = 8 mice from six independent experiments), quinine (medium panel; *n* = 6 mice from four independent experiments), and citric acid (right panel; *n* = 6 mice from four independent experiments). Black circles represent individual measurements from single injections. **(D)** Score for liking reactions during intraoral infusion of water, vehicle solution (0.05% ethanol), and 3.3 µM capsaicin solution (*n*=7 mice from two independent experiments). Black circles represent individual measurements from single injections. **(E)** Score for liking reactions during intraoral infusion of water at room temperature, 50 °C, 70 °C, and 90 °C (*n*=7 mice from two independent experiments). Black circles represent individual measurements from single injections. **(F)** Score for liking reactions during intraoral infusion of water in mice injected with capsazepine or vehicle (*n*=7 mice from two independent experiments). Black circles represent individual measurements from single injections. **(G)** Intake of water and 3.3 µM capsaicin for 18 hours in a two-bottle preference test (*n*=8 mice from two independent experiments). Black circles represent individual measurements from single injections. \**P* < .05, Tukey’s test. W, water; Veh, vehicle; 3.3Cap, 3.3 µM capsaicin; TR, room temperature; CPZ, capsazepine.

The taste reactivity test (Grill and Norgren, 1978; Berridge, 2000) was used to measure the affective reactions of mice in response to intraoral stimulation. The test chamber consisted of a glass or acrylic floor and an acrylic cylinder (30 cm height, 13 cm outside diameter, and 3 mm thickness). A digital video camera (HDR-PJ800; Sony, Tokyo, Japan or HC-VX992MS; Panasonic, Osaka, Japan) was placed beneath the glass floor to record the ventral view of the mouse. On Day 1, mice were placed in the chamber for 15 min for habituation. Every business day from Days 2 to 9, the mice were placed in the chamber and received manual intraoral infusions of 80 µL Milli-Q water for 2 minutes for habituation. Subsequently, all intraoral infusions were performed in a chamber consisting of 80 µL of the solution for 2 minutes. The interval between infusions on the same day was approximately 15 minutes. On Day 10, the mice received an infusion of the 0.44 mM tannic acid. This procedure was repeated twice. On Day 11 (day of tannic acid test), the mice received an infusion of 0.44 mM tannic acid. They then received an infusion of 4.4 mM tannic acid. Subsequently, they received an infusion of 44 mM tannic acid. The tube was washed with Milli-Q water. On Day 12, the mice received an infusion of the 3.3 µM capsaicin. This procedure was repeated twice. On Day 13 (capsaicin test day), the mice received an infusion of 3.3 µM capsaicin. They then received an infusion of 33 µM capsaicin. They then received an infusion of 330 µM capsaicin. The tube was washed with Milli-Q water. On Day 17, the mice received an infusion of the 0.1 mM quinine. This procedure was repeated twice. On Day 18 (quinine test day), the mice received an infusion of 0.1 mM quinine. Subsequently, they were infused with 1 mM quinine. Subsequently, they received an infusion of 10 mM quinine. The tube was washed with Milli-Q water. On Day 19, the mice received an infusion of the 0.01 M citric acid. This procedure was repeated twice. On Day 20 (citric acid test day), the mice received an infusion of 0.01 M citric acid. They then received an infusion of 0.1 M citric acid. Subsequently, they received an infusion of 1 M citric acid. The tube was washed with Milli-Q water. Data from tannic acid stimulation were not analyzed in the present study because we noticed that after completion of the experiments, tannic acid appeared to stimulate not only the somatosensory pathway but also the taste pathway (Soares et al., 2020); thus, it was difficult to interpret that tannic acid-induced reactions are induced by somatosensory stimulation. The reactions of mice during infusions were video recorded for subsequent frame-by-frame video analysis.

### Taste reactivity tests for comparison among water, vehicle, and 3.3 µM capsaicin, and for the assessment of effects of capsazepine and temperature, and a two-bottle preference test

In these tests (Fig. 1D-G; Supplemental fig.1), eight new mice underwent oral surgery, seven mice were used for the taste reactivity test, and all eight mice were used for a two-bottle preference test. One of the eight mice was unable to perform the taste reactivity test because the oral tube was stuck immediately after the surgery. The overall procedure for the taste reactivity test was identical to that for comparison among capsaicin, quinine, and citric acid, as described above with minor modifications. On Day 10, the mice received an infusion of the 3.3 µM capsaicin. This procedure was repeated twice. On Day 11 (test day for comparison between water, vehicle, and 3.3 µM capsaicin), the mice received an infusion of Milli-Q water, vehicle (0.05% ethanol), and 3.3 µM capsaicin. The order in which each solution was presented was counterbalanced between subjects. Immediately after the test of a specific solution, the tube was flashed with Milli-Q water. The interval between infusions on the same day was approximately 30-45 minutes. On Days 12 and 13 (test days for the assessment of the effects of capsazepine), 10 mg/kg BW capsazepine or vehicle was subcutaneously injected 60 min before the taste reactivity tests. For certain mice, capsazepine or vehicle was injected on Day 12, and on Day 13, another solution was injected on Day 12. The order in which each solution was injected was counterbalanced between subjects. The mice received an intraoral infusion of 3.3 µM capsaicin. On Day 14 (test days for the assessment of the effects of temperature), the mice received an infusion of Milli-Q water at room temperature, 50°C, 70°C, and 90°C in order. These temperatures are the temperatures of the water immediately before it is loaded into the syringe. Thus, it is likely that 50 °C, 70 °C, and 90 °C water had somewhat lower temperatures when infused into the oral cavity because hot water can be cooled during transfer through a tube from the syringe to the oral cavity of the mouse. On Days 15-19, a two-bottle preference tests were conducted. All preference tests were conducted in home cages. Fluid was available through a ball-bearing nozzle attached to 10-mL glass bottles (BrainScience idea. Co. Ltd., Tokyo, Japan) that were held on the left or right side of the top of the cage with a spring. Fluid intake was measured to the nearest 0.1 g by weighing the bottles on an electronic balance. habituation (Days 15-17) and tests (Days 18,19) were conducted for 18 hours (17:00-11:00 on the next day). On Day15 (habituation day for the preference test), two bottles containing Milli-Q water were set and measured. On Days 16 and 17 (habituation days for capsaicin in the preference test), two bottles, one containing Milli-Q water and another contained 3.3 µM capsaicin, were set and measured. The left and right positions were exchanged for each subject between Days 16 and 17, and the left-right order was counter-balanced between subjects. On Days 18 and 19 (test days), the same thing was done for Days 16 and 17 and was the result of the preference test.

### Taste reactivity video scoring

Manual video analysis was performed as previously described (Tanaka et al., 2019) with minor modifications. Liking reactions included midline or lateral tongue protrusion (emergence of the anterior tip of the tongue, which pushes both the left and right upper lips or either the left or right lip, respectively), and paw licking (licking with tongue emergence toward the mouse’s forepaws, with the paws held close to the mouth). The number of tongue protrusions was quantified as the sum of midline and lateral tongue protrusions.

Disgust reactions included gape (large opening of the mouth with retraction of the lower lip), headshake (rapid lateral movement of the head), face wash (wipes over the face with the paws), forelimb flail (rapid waving of both forelimbs), and floor and wall chin rubs (pushing the chin against the floor or wall of the test chamber, respectively). The number of chin rubs was quantified as the sum of the number of floor and wall chin rubs. Neutral reactions (less strongly liked to liking/disgust evaluations) were considered rhythmic mouth movements (mouth movements without the emergence of the tongue) and ordinary grooming (sequential movements of behavior that include movements that bring the mouth close to the abdomen or genital area). To ensure that each component of taste reactivity contributed equally to the final scores, reactions that occurred in continuous bouts were scored in time bins (Berridge 2000) with minor modifications. Components characterized by long-duration bouts, such as paw licking, face wash, mouth movements, and grooming, were scored in five-second bins (successive repetitions within five seconds were scored as one occurrence). Components characterized by moderate-duration bouts, such as chin rub were scored in two-second bins. Other reactions, such as tongue protrusion and forelimb flail, which can occur as a single reaction, were scored as separate occurrences (e.g., one forelimb flail equals one occurrence). The total liking reaction was quantified as the sum of the scores for tongue protrusion and paw licking. The total disgust reaction was quantified as the sum of the scores for gape, headshake, face wash, forelimb flail, and chin rub. The total neutral reaction was quantified as the sum of the scores for mouth movement and grooming.

### Statistical analysis

Differences between more than three groups were analyzed using repeated-measures one-way analysis of variance (ANOVA) followed by Tukey’s multiple comparison test (Supplemental table 1). The differences between the two groups were analyzed using the Wilcoxon test. Statistical significance was set at P < 0.05. All tests were two tailed. Most data are expressed as line graphs with dots representing the group mean and error bars representing the standard error of the mean (SEM), or truncated violin plots with gray bars representing the group median and dotted lines representing the interquartile range. Numerical data represented in the main text are presented as mean ± SEM. The comprehensive and detailed results of the statistical analyses are summarized in Supplementary table 1.

## Results

To investigate the orofacial and bodily reactions induced by intraoral infusion of capsaicin and compare them with those induced by quinine and citric acid, three different concentration solutions were infused for each stimulus. The sum of the scores for tongue protrusion and paw licking was assessed as the total liking reaction (Berridge, 2000). The sum of the scores of gape, headshake, face wash, forelimb flail, and chin rub was assessed as the total disgust reaction.

The low (3.3 µM)-and medium (33 µM)-concentration capsaicin induced liking reactions (6.7 ± 1.8 scores/ minute at 3.3 µM; 5.8 ± 1.7 at 33 µM; *n* = 8 mice from six independent experiments; Fig. 1A-C), which are comparable to those induced by the sweet saccharin solution in our previous study (6.9 ± 1.6; Tanaka et al., 2019). The 3.3 µM capsaicin induced significantly more liking reactions than water or vehicle (0.05% ethanol) solution (*n*=7 mice from two independent experiments; Fig. 1D, Supplemental fig.1A,B). Since the capsaicin receptor, TRPV1, is a heat receptor and liking reactions include tongue protrusions, one may argue that capsaicin makes mice sense heat on their tongues and mice may extend their tongues to cool them down, resulting in increased liking reactions. To test this possibility, we infused hot water and found that it did not induce significant liking reactions (*n*=7 mice from two independent experiments; Fig. 1E, Supplemental fig.1C,D), suggesting that heat signaling, which might be activated by capsaicin, was not sufficient to induce liking reactions. The capsaicin-induced liking reactions were significantly suppressed by pretreatment with capsazepine, an inhibitor of the TRPV1 channel (*n*=7 mice from two independent experiments; Fig. 1F, Supplemental fig.1E,F), suggesting that capsaicin induced liking reactions through the TRPV1 channel, at least in part. Consistent with previous studies (Bachmanov et al., 1996; Simons et al., 2001; Furuse et al., 2002; Narukawa and Misaka, 2021), 3.3 µM capsaicin was avoided in a two-bottle preference test compared with water (*n*=8 mice from two independent experiments; Fig 1G). Thus, 3.3 µM capsaicin induced significantly more liking reactions (Fig. 1D), but was significantly avoided (Fig.1G). A weaker but similar elicitation of liking reactions was observed with citric acid (*n* = 6 mice from four independent experiments; Fig. 1B, C), which is consistent with previous studies (Brining et al., 1991; Galaverna et al., 1993; Steiner, 2001). All quinine concentrations induced almost no liking reactions (*n* = 6 mice from four independent experiments; Fig. 1B, C). Capsaicin-induced liking reactions were not almost completely induced at the highest concentration (*n* = 8 mice from six independent experiments; 0.1 ± 0.1 scores/ minute; Fig. 1A, B). At the lowest concentrations for each stimulus, all of three stimuli induced few disgust reactions (1.9 ± 0.7 at 3.3 µM capsaicin, *n* = 8 mice from six independent experiments; 4.8 ± 1.9 at 10^2^ µM quinine, *n* = 6 mice from four independent experiments; 2.9 ± 0.6 at 10^4^ µM citric acid, *n* = 6 mice from four independent experiments; Fig. 2), which are comparable to those induced by the water infusion (2.1 ± 1.6, *n*=7 mice from two independent experiments; Supplemental fig.1B). At the highest concentrations, all three stimuli induced significantly higher scores for disgust reactions than those at lower concentrations (*n* = 8 mice from six independent experiments for capsaicin, *n* = 6 mice from four independent experiments for quinine and citric acid; Fig. 2; Supplemental fig.2). Thus, capsaicin elicits both liking and disgust reactions, depending on its concentration.

**Figure 2.**
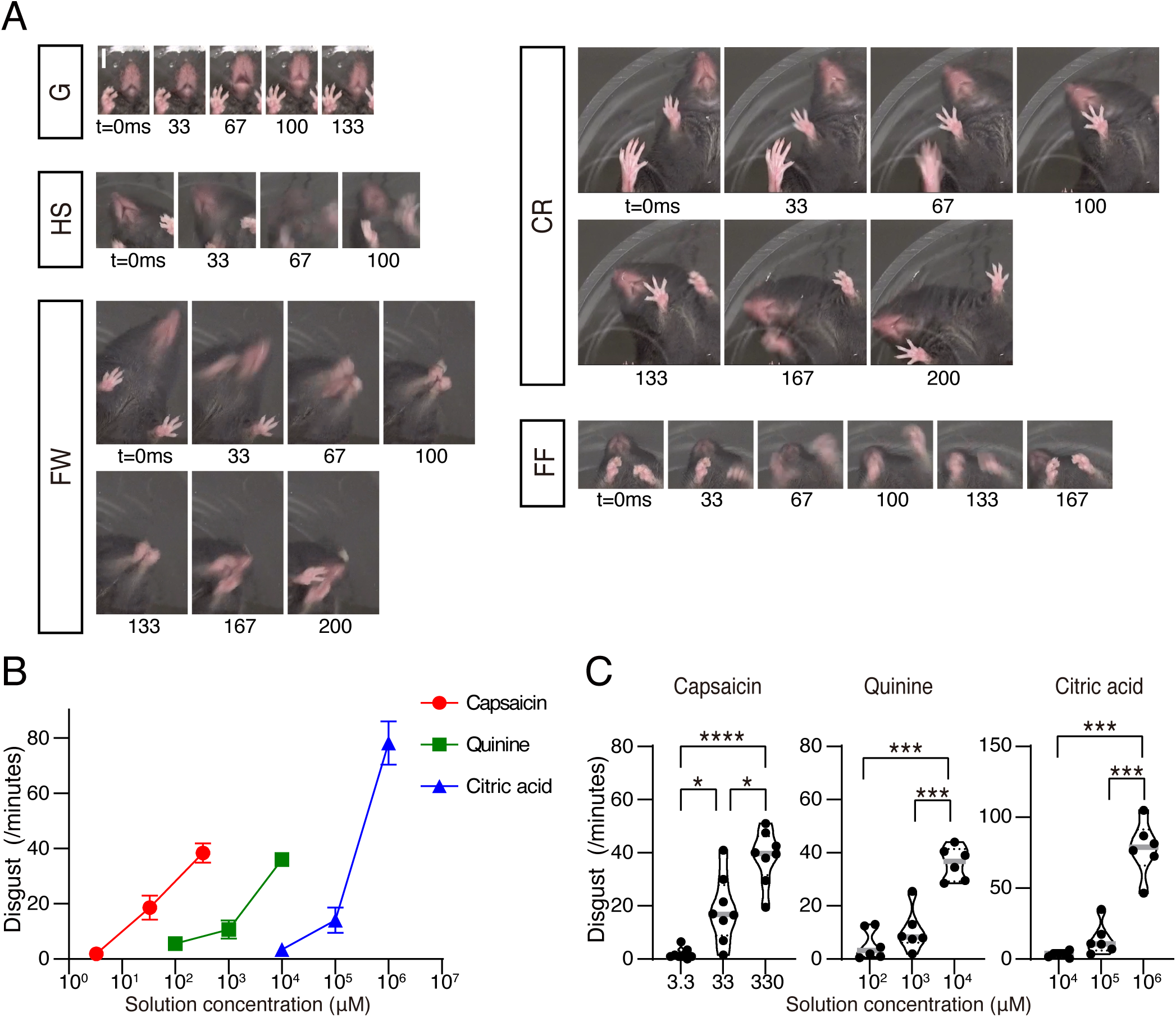
Capsaicin of increasing concentration elicits more disgust reactions. **(A)** Representative time-lapse sequences of disgust reactions during the intraoral infusion of capsaicin. The numbers at the bottom of each panel indicate the elapsed time (milliseconds). G, gape; HS, head shake; FW, face wash; CR, chin rub; FF, forelimb flail. Scale bar, 5 mm. **(B)** Relationship between the score of disgust reactions during intraoral infusion of capsaicin (red circles; *n* = 8 mice from 6 independent experiments), quinine (green rectangles; *n* = 6 mice from 4 independent experiments), and citric acid (blue triangles; *n* = 6 mice from 4 independent experiments) and their concentrations. Error bars are SEM. **(C)** The score of disgust reactions during intraoral infusion of three different concentrations of capsaicin (left panel; *n* = 8 mice from six independent experiments), quinine (medium panel; *n* = 6 mice from four independent experiments), and citric acid (right panel; *n* = 6 mice from four independent experiments). Black circles represent individual measurements from single injections. \**P* < .05, \*\*\**P* < .001, \*\*\*\**P* < .0001, Tukey’s test.

As the concentration of capsaicin increased, disgust reactions increased and liking reactions decreased as a whole (*n* = 8 mice from six independent experiments; Fig.3A). A similar trend was observed for citric acid (*n* = 6 mice from four independent experiments; Fig.3A). The mechanistic relationship between the liking and disgust reactions could potentially be one of the following (Fig. 3B). First, the liking reactions decrease as the disgust reactions increase in a seesaw-like manner. In this *seesaw* model, the increase in disgust reactions is always accompanied by a decrease in liking reactions. Second, the liking reactions change independently of the disgust reactions. In this *independent* model, when disgust reactions increase, liking reactions may also increase. We analyzed how the disgust and liking reactions of each mouse changed as the concentration of capsaicin or citric acid solution increased from low to medium concentrations and from medium to high concentrations. With respect to changes in capsaicin (*n* = 8 mice from six independent experiments; Fig. 3C) and citric acid (*n* = 6 mice from four independent experiments; Fig. 3D) concentrations from low to medium, all mice showed an increase in disgust reactions, and approximately half of them also showed an increase in liking reactions, which is consistent with an *independent* model. With respect to changes in capsaicin (*n* = 8 mice from six independent experiments; Fig. 3E) and citric acid (*n* = 6 mice from four independent experiments; Fig. 3F) concentrations from medium to high, most mice showed an increase in disgust reactions, and most of them showed a decrease in liking reactions, consistent with a *seesaw* model. These data suggest that the mechanistic relationship between liking reactions and disgust reactions is dynamically regulated, depending on the intensity of the chemosomatosensory and taste stimuli that induce them.

**Figure 3.**
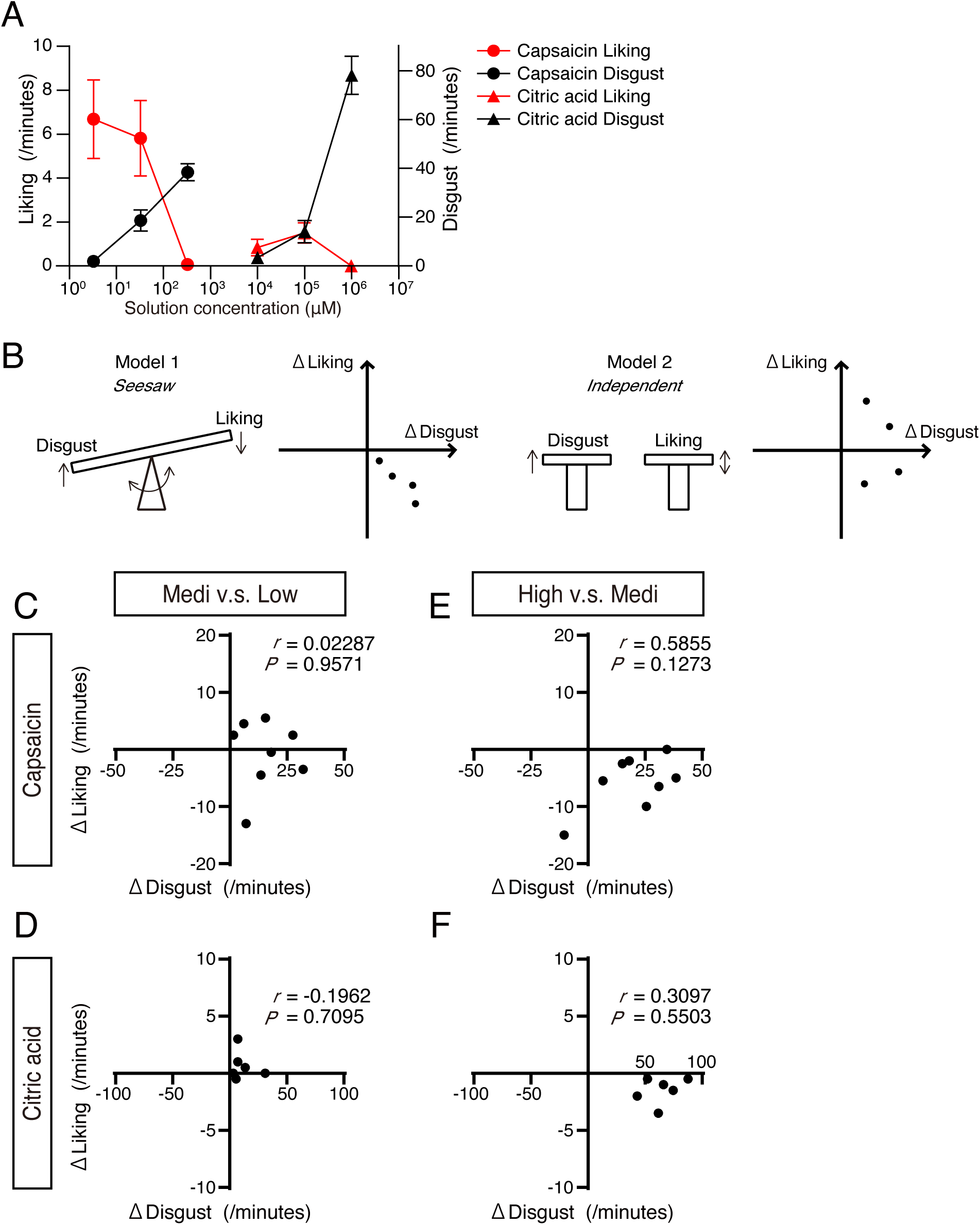
The mechanistic relationship between liking reactions and disgust reactions is dynamically regulated depending on the solution concentration. **(A)** Relationship between liking reaction scores (red) and disgust reactions (black) during intraoral infusion of capsaicin (circles) and citric acid (triangles), and their concentrations. Graphs showing excerpts of the data shown in Fig. 1B and Fig. 2B are overlaid on top of each other. **(B)** Diagram of possible models explaining the mechanistic relationship between liking and disgust reactions. In the *seesaw* model, an increase in disgust reactions is always accompanied by a decrease in liking reactions. In the *independent* model, when disgust reactions increase, liking reactions may also increase. **(C)** The amount of change from the scores of liking and disgust reactions induced by medium-concentration (33 µM) capsaicin to those induced by low-concentration (3.3 µM) capsaicin in each mouse. Black circles represent the difference between two individual measurements from a single injection in each mouse. **(D)** The amount of change from the scores of liking and disgust reactions induced by medium-concentration (0.1 M) citric acid to those induced by low-concentration (0.01 M) citric acid in each mouse. Black circles represent the difference between two individual measurements from a single injection in each mouse. **(E)** The amount of change from the scores of liking and disgust reactions induced by high-concentration (330 µM) capsaicin to those induced by medium-concentration (33 µM) capsaicin in each mouse. Black circles represent the difference between two individual measurements from a single injection in each mouse. **(D)** The amount of change from the scores of liking and disgust reactions induced by high-concentration (1 M) citric acid to those induced by medium-concentration (0.1 M) citric acid in each mouse. Black circles represent the difference between two individual measurements from a single injection in each mouse. *r*, Pearson’s *r*.

Previous studies suggest that two distinct criteria must be satisfied for liking and disgust reactions to be considered indicators of liking and disgust in animals (Berridge, 2000). First, multiple orofacial and bodily reactions that comprise each of the liking and disgust reactions should show a similar direction of change (increase or decrease) (Fig. 4A, B). To test whether the reactions induced by capsaicin meet this first criterion, as do those induced by taste stimuli, we systemically analyzed each reaction. As for liking reactions, defined as the sum of scores of tongue protrusion and paw licking, only tongue protrusion was observed (Fig.4C, Supplemental fig. 3A), and paw licking was not observed in any of the tests in any mice (*n* = 8 mice from six independent experiments for capsaicin, *n* = 6 mice from four independent experiments for quinine and citric acid). In contrast, medium-and high-concentration capsaicin increased various disgust reactions as well as high-concentration quinine and citric acid in a concentration-dependent manner (Fig. 4D, Supplemental fig. 3B), although head shake was rarely observed under any of the tested conditions (*n* = 8 mice from six independent experiments for capsaicin, *n* = 6 mice from four independent experiments for quinine and citric acid). Thus, disgust reactions induced by medium-and high-concentration capsaicin satisfied the first criterion, as well as those induced by high-concentration quinine and citric acid. Second, when the liking or disgust reaction changes, another reaction and a neutral reaction must remain unchanged or exhibit opposite directional changes (Fig. 5A, B). As for liking reactions, low-and medium-concentration capsaicin increased liking reactions compared to high-concentration capsaicin, and simultaneously decreased disgust reactions (*n* = 8 mice from six independent experiments; Fig. 5C). However, low and medium concentrations of capsaicin increased neutral reactions (*n* = 8 mice from six independent experiments; Fig. 5C, Supplemental fig. 4A, B). In contrast, high-concentration capsaicin increased disgust reactions and simultaneously decreased both liking reactions and neutral reactions (*n* = 8 mice from six independent experiments; Fig. 5C, Supplemental fig. 4A, B). The medium concentration of capsaicin increased disgust reactions and did not significantly change either liking or neutral reactions (*n* = 8 mice from six independent experiments; Fig. 5C, Supplemental fig. 4A, B). High-concentration quinine (*n* = 6 mice from four independent experiments; Fig. 5D) and citric acid (*n* = 6 mice from four independent experiments; Fig. 5E) induced largely similar reactions to medium-and high-concentration capsaicin (*n* = 8 mice from six independent experiments; Fig. 5C). For each reaction categorized as a neutral reaction, mouth movements were significantly decreased by high-concentration capsaicin compared to low-and medium-concentration capsaicin (Supplemental fig. 4C, D), while grooming was never observed in all tests in any mice (*n* = 8 mice from six independent experiments for capsaicin, *n* = 6 mice from four independent experiments for quinine and citric acid). Thus, disgust reactions induced by medium-and high-concentration capsaicin satisfied the second criterion, as well as those induced by high-concentration quinine and citric acid. Taken together, while liking reactions induced by low-and medium-concentration capsaicin pass neither the first nor the second criterion, disgust reactions induced by medium-and high-concentration capsaicin satisfy both the first and second criteria, as well as those induced by high-concentration quinine and citric acid.

**Figure 4.**
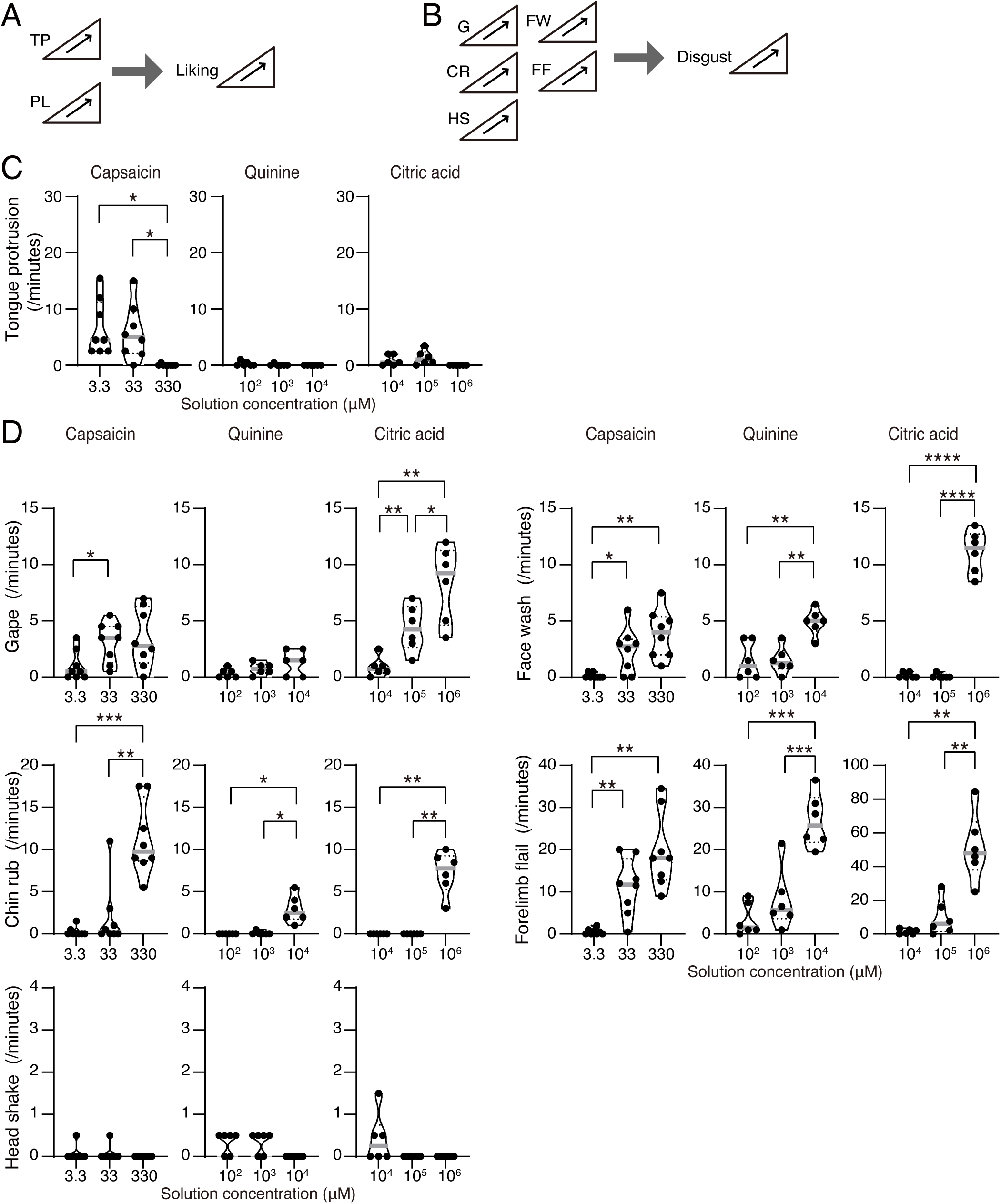
Various reactions consisting capsaicin-induced disgust reactions show a similar direction of change as the increase of capsaicin concentration. **(A, B)** Diagrams explaining that multiple orofacial and somatic reactions that comprise each of the liking (A) and disgust (B) reactions should show a similar direction of change. TP, tongue protrusion; PL, paw licking; G, gape; CR, chin rub; HS, head shake; FW, face wash; FF, forelimb flail. **(C, D)** The scores of tongue protrusion (**C**), gape, chin rub, head shake, face wash, and forelimb flail (**D**) during intraoral infusion of three different concentrations of capsaicin (*n* = 8 mice from six independent experiments), quinine (*n* = 6 mice from four independent experiments), and citric acid (*n* = 6 mice from four independent experiments). Black circles represent individual measurements from single injections. \**P* < .05, \*\**P* < .01, \*\*\**P* < .001, \*\*\*\**P* < .0001, Tukey’s test.

**Figure 5.**
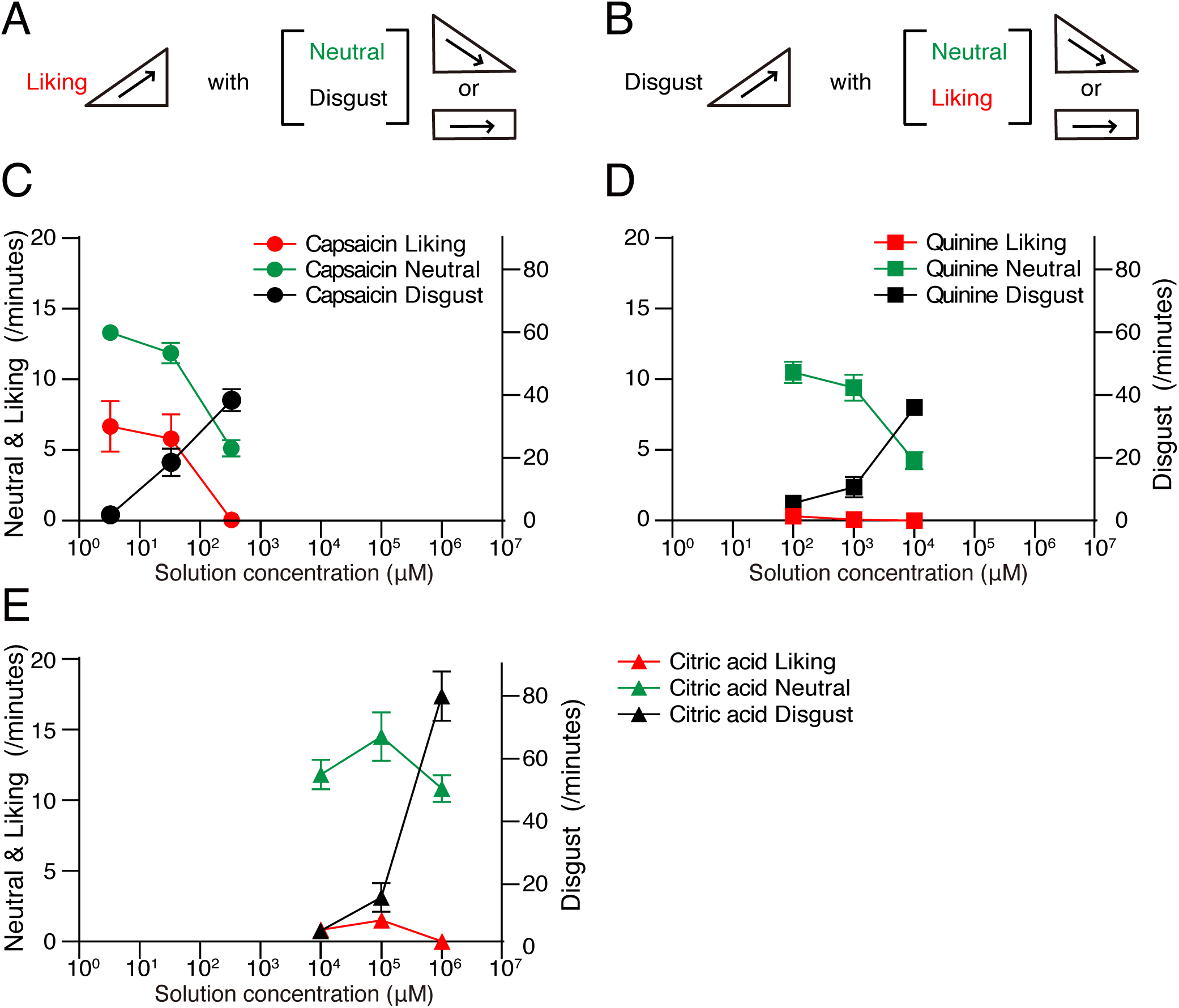
Increase of capsaicin-induced disgust reactions accompanies with decrease or constant of liking reactions and neutral reactions. **(A, B)** Diagrams explaining that when the liking reaction (**A**) or disgust reaction (**B**) increases, another reaction and a neutral reaction should remain unchanged or change in the opposite direction. **(C-E)** Relationship between the scores of the liking reaction (red), neutral reaction (green), and disgust reaction (black) during intraoral infusion of capsaicin (circles), quinine (rectangle), and citric acid (triangles), and their concentrations. Graphs showing excerpts of capsaicin (**C**), quinine (**D**), and citric acid (**E**) data shown in Fig. 1B, Fig. 2B, and Supplemental fig. 3A, are overlaid on top of each other.

Previous studies suggest that as the total disgust score was increased, the ratio of each disgust reaction among all disgust reactions changes: gape appeared first at the lowest disgust score in total, then chin rub was emitted as the total disgust score increased, and finally, forelimb flail appeared (Breslin et al., 1992). To determine whether similar changes in the reaction were observed with capsaicin stimulation, we analyzed the ratio of each disgust reaction among all disgust reactions at different concentrations of capsaicin and compared them with those induced by quinine and citric acid. As the concentration of the solution was increased, the ratio of each disgust reaction changed similarly among capsaicin, quinine, and citric acid; gape and headshake were observed from the beginning and tended to decrease as the concentration of the solution increased (Fig. 6A, C), face wash, and forelimb flail were consistently observed at all concentrations (Fig. 6D, E), and the ratio of chin rub significantly increased at high concentrations (*n* = 8 mice from six independent experiments for capsaicin, *n* = 6 mice from four independent experiments for quinine and citric acid; Fig. 6B, F, Supplemental fig. 5). These results indicate that the changes in the ratio of each disgust reaction as a function of the intensity of the capsaicin stimulus are similar to those in response to quinine and citric acid. These findings suggest that the components of disgust reactions are dynamically regulated depending on the intensity of the chemosomatosensory and taste stimuli that induce them.

**Figure 6.**
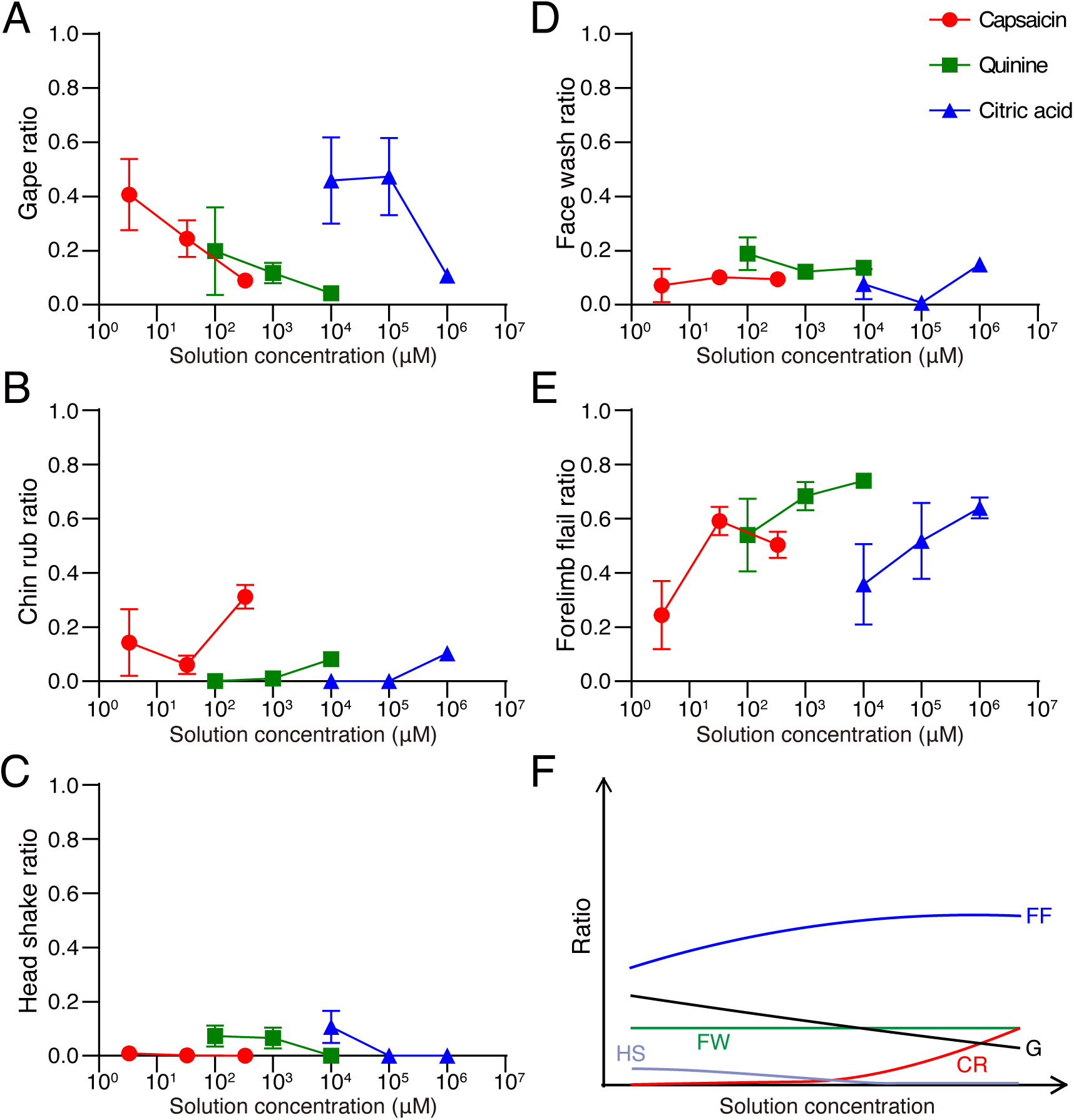
Capsaicin induces various disgust reactions in a concentration-dependent manner. **(A-E)** The ratio of gape (**A**), chin rub (**B**), head shake (**C**), face wash (**D**), and forelimb flail **(E)** among all disgust reactions induced by capsaicin (*n* = 8 mice from six independent experiments), quinine (*n* = 6 mice from four independent experiments), and citric acid (*n* = 6 mice from four independent experiments). **(F)** Illustration summarizing the results of **A-E**. Gape and headshake were observed from the beginning, and tended to decrease as the concentration of the solution increased. Face wash and forelimb flail were constantly observed at all concentrations. The chin rub increased at high concentrations of capsaicin, quinine, and citric acid. Error bars are SEM. G, gape; CR, chin rub; HS, head shake; FW, face wash; FF, forelimb flail.

## Discussion

The present study demonstrated that intraoral capsaicin stimulation elicited liking reactions at low concentration (Fig. 1) and disgust reactions at high concentration (Fig. 2) in mice. This is the first demonstration of non-taste stimuli eliciting liking and disgust reactions, defined as taste reactivity (Grill and Norgren, 1978; Berridge, 2000). Given that liking and disgust reactions reflect biological processes, which may underlie human emotions of liking and disgust (Berridge, 2000), the present results indicate the possibility that liking and disgust reactions can be used as a measure to quantitatively estimate subjective liking and disgust, irrespective of stimulation modalities in mice.

### Technical consideration

The primary aim of the present study was to determine whether stimuli from modalities other than taste, such as capsaicin, could induce behaviors defined in taste reactivity tests. Previous studies have suggested that after exposure to taste stimuli, taste reactivity in the following test may be influenced by prior stimuli (Pecina and Berridge, 2005; Ho and Berridge, 2014; Castro and Berridge, 2017). To eliminate the possibility that taste reactivity to capsaicin might be altered by the influence of taste stimuli, such as high-concentration quinine solutions, we conducted taste reactivity tests in the order of capsaicin, quinine, and citric acid. However, the disgust reactions for citric acid presented in the present study might have been influenced by prior quinine stimulation due to this testing sequence. Another possibility is that mice exposed to high concentrations of capsaicin and/or quinine entered a state of learned helplessness, which could result in reduced disgust reactions to quinine and/or citric acid. Nonetheless, since the primary objective of this study was not to compare the responses to capsaicin, quinine, and citric acid, we believe that the effects of these possibilities on the main conclusions of the present study are limited.

### Mice may like low-concentration capsaicin as well as humans

The present study demonstrated that intraoral low concentration (3.3 µM) capsaicin induced liking reactions (Fig. 1A-D) but not the disgust reactions (Fig. 2B,C) in taste reactivity tests. It is believed that only humans like capsaicin, and other animals do not. Capsaicin is probably the most widely consumed spice in the world (Rozin and Schiller, 1980), and intraoral 3-13.2 µM capsaicin induces liking in humans with large individual differences (Rozin and Schiller, 1980; Rozin et al., 1982; Stevenson and Yeomans, 1993; 1995; Tornwall et al., 2012; Byrnes and Hayes, 2016). In contrast, animals consistently avoid capsaicin (Bachmanov et al, 1996; Simons et al., 2001; Furuse et al., 2002; Narukawa and Misaka, 2021). In the present study, low-concentration (3.3 µM) capsaicin was indeed avoided in the two-bottle preference test (Fig. 1G). Thus, the mice showed sole liking reactions (Fig. 1A-D) but not the disgust reactions (Fig. 2B,C) to 3.3 µM capsaicin in the taste reactivity test, but avoided them in the two-bottle preference test (Fig. 1G). These seemingly conflicting results are interesting but not surprising because it is well known that liking reactions in taste reactivity reflect a hedonic core process (Berridge, 2000) but not intake preference measured in choice tests (Bachmanov et al., 1996; Simons et al., 2001; Furuse et al., 2002; Narukawa and Misaka, 2021). Thus, 3.3 µM capsaicin activated the core hedonic process but did not activate the preference process. Stimuli, such as sucrose, which elicit liking reactions, are generally preferred. To the best of our knowledge, 3.3µM capsaicin is the first stimulus demonstrated to trigger a liking reaction while being avoided.

Both capsaicin-induced and citric acid-induced liking reactions were completely dominated by tongue protrusions (compare Fig. 1C with Fig. 4C). Chemical irritants, such as formalin, mustard oil, and capsaicin, injected into the paw often induce licking, biting, and shaking of the paw in rodents (Koepp et al., 2006; Kwan et al., 2006; Sakurada et al., 2009). One may argue that capsaicin-induced tongue protrusions reflect licking behavior against chemical nociception. Of note, the highest concentration (330 µM) of capsaicin, which is much higher than the concentration that induces intraoral pain in humans (30 µM) (Ngom et al., 2001), did not induce tongue protrusion (Fig. 4C), but induced disgust (Fig.2C), suggesting that painful capsaicin stimulus in the oral cavity does not necessarily induce tongue protrusion. Thus, tongue protrusions induced by intraoral low-concentration capsaicin likely represent liking, but not licking against chemical nociception.

Taken together, the present results support the notion that mice like 3.3 µM capsaicin as well as humans. To complement the evolutionary gap between liking reactions in mice in the present study and conscious liking ratings in humans (Rozin and Schiller, 1980; Rozin et al., 1982; Stevenson and Yeomans, 1993; 1995; Tornwall et al., 2012; Byrnes and Hayes, 2016), it would be interesting to test whether not only sweet taste (Steiner, 2001) but also 3.3 µM capsaicin induces liking reactions in human neonates and monkeys.

Individual differences in liking for capsaicin in humans may be explained, at least in part, by differences in sensitivity to capsaicin. Capsaicin non-likers tend to report higher sensory intensity ratings than likers do (Stevenson and Yeomans, 1993; Tornwall et al., 2012). However, even with the same level of perceived intensity, affective responses differ significantly between likers and non-likers (Stevenson and Yeomans, 1993). This suggests that factors other than sensitivity may also play a role, although the factors remain unclear. In the present study, we assessed the degree of liking for capsaicin in an animal model. This experimental setup is expected to help explore the mechanisms underlying the liking of capsaicin in future research.

### Mice may experience disgust from intraoral medium-and high-concentration capsaicin

While 3.3 µM capsaicin induced few disgust reactions (Fig. 2B, C) comparable to water (2.4 ± 1.0; Tanaka et al., 2019), 33 µM and 330 µM capsaicin induced significant disgust reactions (Fig. 2 B, C). 33 µM capsaicin also induced significant liking reactions, suggesting that it induced both liking and disgust, according to the two-dimensional account of palatability (Berridge and Grill, 1983; 1984). Given that 33 µM and 330 µM capsaicin are higher than 12-13.2 µM capsaicin that induced unpleasantness or disliking in humans with large individual differences (Stevenson and Yeomans, 1993; Byrnes and Hayes, 2016), 33 µM and 330 µM capsaicin infusion may induce not only disgust reactions but also unpleasantness, disliking, and/or disgust in mice. To test this hypothesis, it is crucial to test whether intraoral 33 µM or 330 µM capsaicin induces disgust in humans. In addition, to complement the evolutionary gap between disgust reactions in mice and conscious disgust ratings in humans, it would be interesting to test whether not only bitter taste (Steiner et al., 2001), but also 33 µM or 330 µM capsaicin induces disgust reactions in human neonates and monkeys.

### The potential interaction of chemesthesis with taste in emotions

Previous studies have shown that suprathreshold concentrations of capsaicin diminish the intensity of sweet and bitter tastes (Prescott and Stevenson, 1995; Simons et al., 2002; Kapaun and Dando, 2017). Conversely, peri-threshold concentrations (∼1µM) of capsaicin enhance sensitivity to sweet, bitter, sour, and umami tastes (Narukawa et al., 2011; Han et al., 2022). Additionally, sweetness has been found to reduce the spiciness intensity of capsaicin (Smutzer et al., 2018). Our study revealed that chemical somatosensory stimulation with capsaicin and gustatory stimulation can elicit a variety of similar orofacial responses depending on the concentration of the stimulus. In the future, investigating how these orofacial responses change due to the interaction between chemical somatosensory and gustatory stimuli could lead to a better understanding of the neural mechanisms underlying the emotions generated by these interactions.

### Intraoral medium-and high-concentration capsaicin may induce not only disgust reactions but also pain in mice

In the present study, 33 µM and 330 µM capsaicin induced disgust reactions (Fig. 2B, C). Given that 33 µM and 330 µM capsaicin are in the range of 30 µM-165 mM capsaicin that induces intraoral pain in humans (Ngom et al., 2001; Baad-Hansen et al., 2003; Zhao et al., 2012; Lu et al., 2013; Naganawa et al., 2015), intraoral 33 µM and 330 µM capsaicin are likely to induce not only disgust reactions but also pain in mice. Although the relationship between disgust and pain remains unclear, they can be tightly linked: disgust and pain share some facial action units (Kappesser and Williams, 2002; Kunz et al., 2013) and neuronal responses in humans (Corradi-Dell’Acqua et al., 2016). It would be interesting to investigate the similarities and differences in the underlying neural mechanisms of disgust and pain.

### Capsaicin-induced disgust reactions to investigate the mechanism of oral disorders in mice

Assessing disgust and pain levels arising from the oral cavity in an awake animal model is essential for elucidating the pathophysiology of oral discomfort and pain syndromes, including oral cenesthopathy (Umezaki et al., 2016) and burning mouth syndrome (Klein et al., 2020). However, no awake-animal model is available to date, probably because experimental manipulation of intraoral tissues is challenging in awake animals (Terayama et al., 2014). The present study utilized a taste reactivity procedure developed in the research field of taste and emotion (Grill and Norgren, 1978; Berridge, 2000) to stimulate intraoral tissue with high-concentration capsaicin, which induces unpleasantness (Stevenson and Yeomans, 1993; Byrnes and Hayes, 2016) and pain (Ngom et al., 2001; Baad-Hansen et al., 2003; Zhao et al., 2012; Lu et al., 2013; Naganawa et al., 2015) in humans, and detected disgust reactions induced by stimulation (Fig. 2B, C). Thus, capsaicin-induced disgust reactions may be used as a readout of intraoral discomfort and pain in mice to elucidate the mechanisms underlying the pathophysiology of oral discomfort and pain syndromes (Umezaki et al., 2016; Klein et al., 2020). As the first step in assessing this assumption, it would be interesting to test whether effective drugs for oral cenesthopathy, such as aripiprazole (Umezaki et al., 2018), and for burning mouth syndrome, such as clonazepam (Klein et al., 2020), suppress capsaicin-induced disgust reactions.

## Acknowledgments

We thank Mr. Shizuki Inaba and other lab members for their helpful comments and discussion.

## Data Availability

The data underlying this article will be shared upon reasonable request to the corresponding author.

