## Supplemental figures, table and legends for "Innate liking and disgust reactions elicited by intraoral capsaicin in male mice"

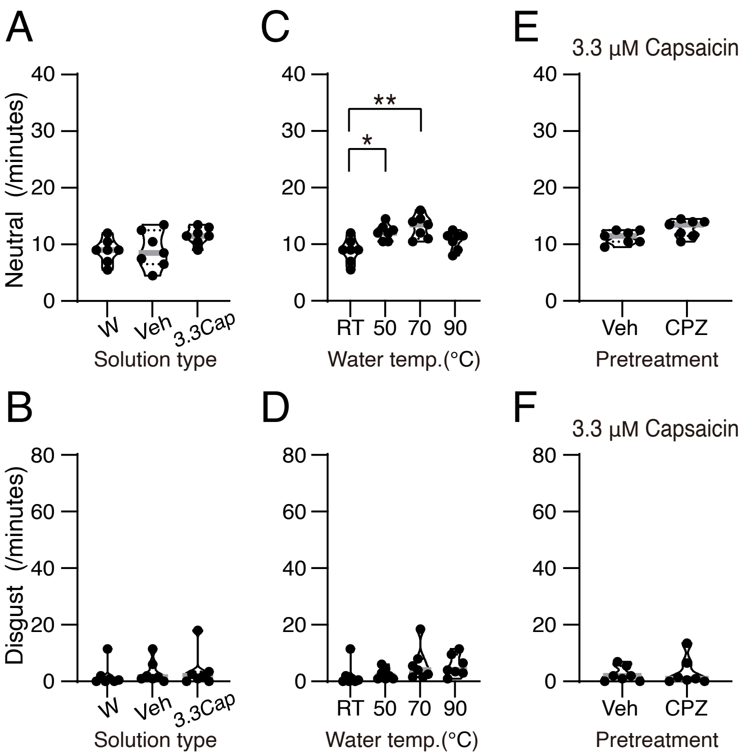


**Supplemental figure 1**

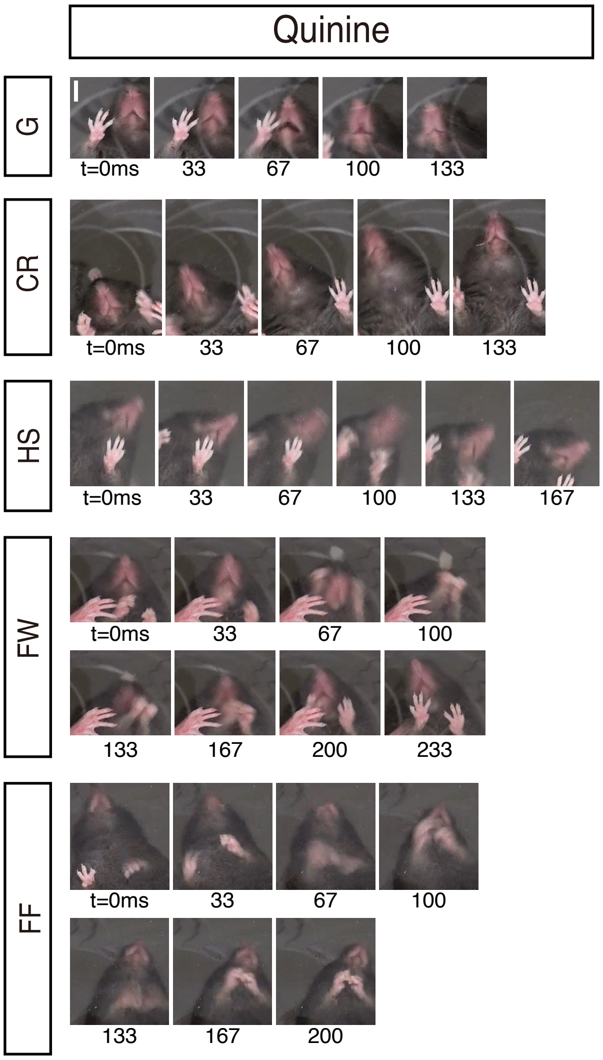


**Supplemental figure 2**

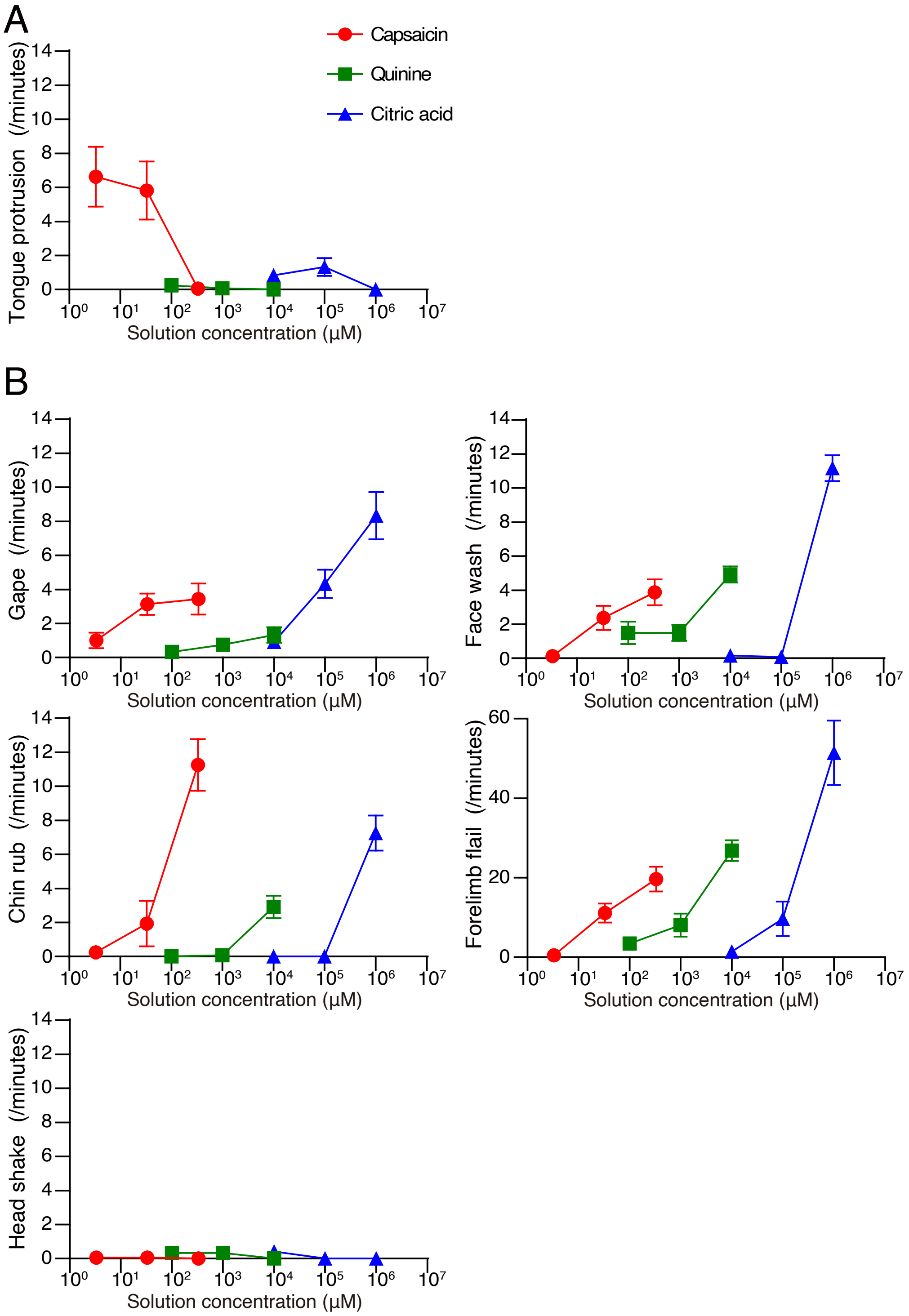


**Supplemental figure 3**

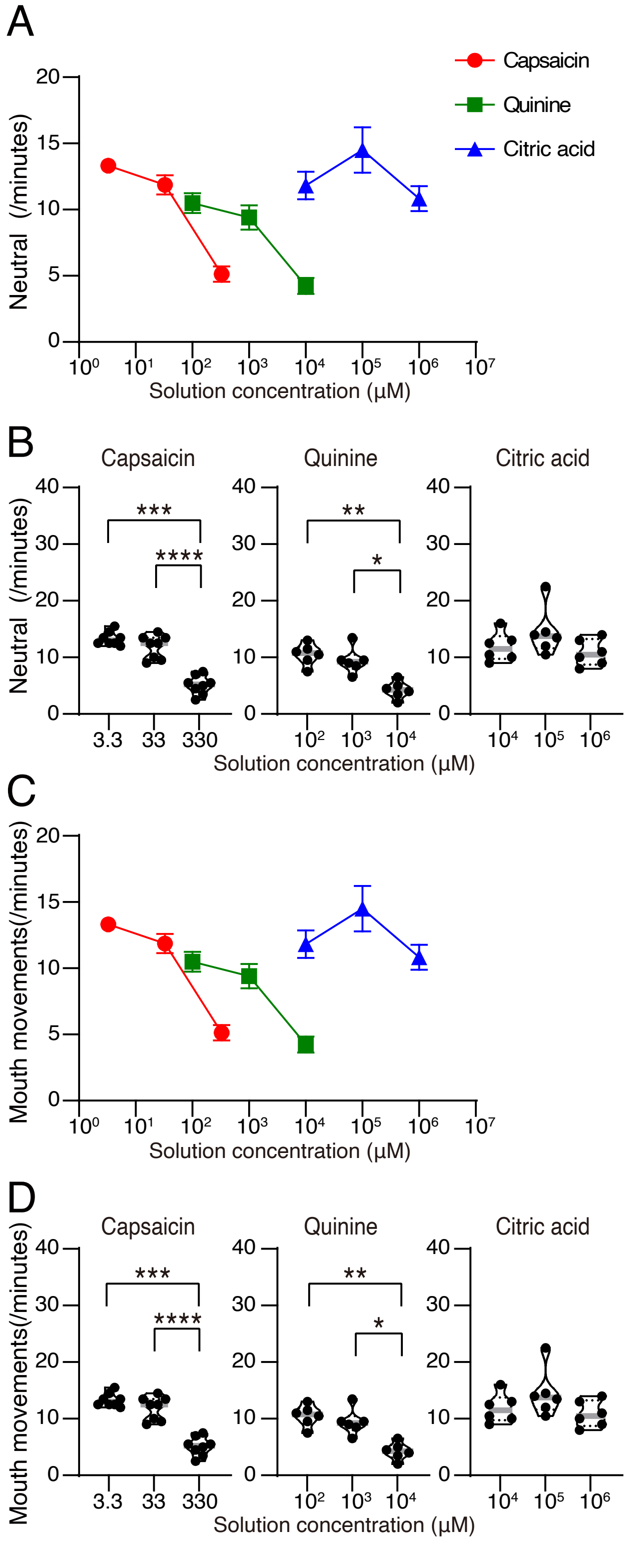


**Supplemental figure 4**

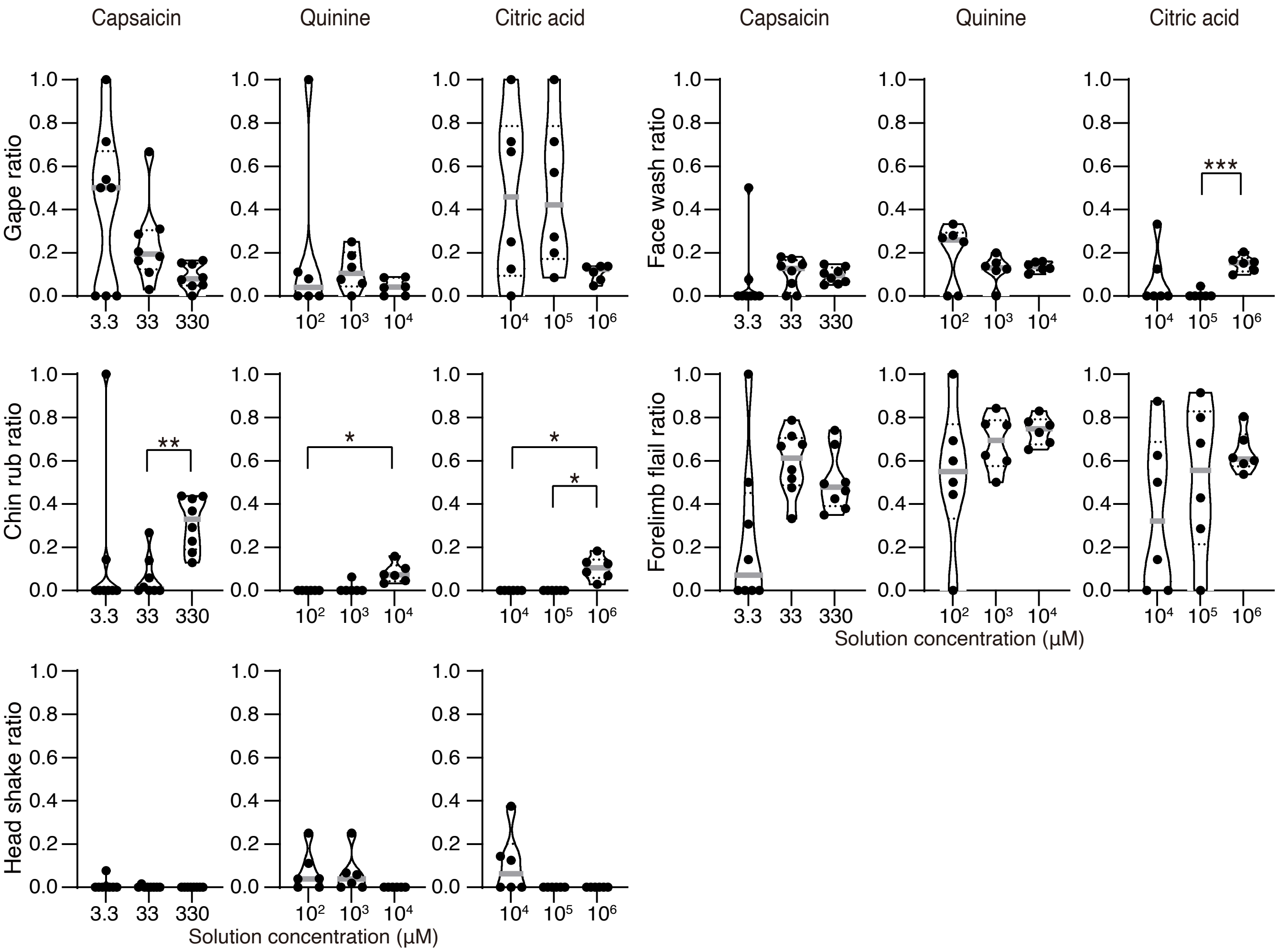


**Supplemental figure 5**

**Supplemental table 1** Han et al.

| | | | | | | |
|---|---|---|---|---|---|---|
| Fig.1C | Liking Capsaicin | RM one-way ANOVA | 3.3μL vs. 33μL | Tukey's | ns | 0.913 |
| | | F (1.854, 12.98) = 7.215 | 3.3μL vs. 330μL | Tukey's | * | 0.0193 |
| | | **P=0.0087 | 33μL vs. 330μL | Tukey's | * | 0.0284 |
| | Liking Quinine | RM one-way ANOVA | 100μL vs. 1000μL | Tukey's | ns | 0.6083 |
| | | F (1.000, 5.000) = 1.000 | 100μL vs. 10000μL | Tukey's | ns | 0.6083 |
| | | P=0.3632 | 1000μL vs. 10000μL | Tukey's | ns | 0.6083 |
| | Liking Citric acid | RM one-way ANOVA | 10000μL vs. 100000μL | Tukey's | ns | 0.4519 |
| | | F (1.806, 9.030) = 5.446 | 10000μL vs. 1000000μL | Tukey's | ns | 0.1655 |
| | | *P=0.0303 | 100000μL vs. 1000000μL | Tukey's | ns | 0.0517 |
| Fig.1D | Liking solution type | RM one-way ANOVA | Water vs. 0.05%EtOH | Tukey's | ns | 0.9168 |
| | | F (1.155, 6.931) = 10.78 | Water vs. 3.3 | Tukey's | * | 0.0288 |
| | | *P=0.0121 | 0.05%EtOH vs. 3.3 | Tukey's | * | 0.0417 |
| Fig.1E | Liking Hot water | RM one-way ANOVA | Water vs. 50°C | Tukey's | ns | 0.9977 |
| | | F (1.601, 9.606) = 0.4167 | Water vs. 70°C | Tukey's | ns | 0.9374 |
| | | P=0.6274 | Water vs. 90°C | Tukey's | ns | 0.9549 |
| | | | 50°C vs. 70°C | Tukey's | ns | 0.5773 |
| | | | 50°C vs. 90°C | Tukey's | ns | 0.7554 |
| | | | 70°C vs. 90°C | Tukey's | ns | 0.9438 |
| Fig.1F | Liking | CPZ vs. DMSO | Wilcoxon test | two-tailed | * | 0.0313 |
| Fig.1G | Two-bottle test | Water vs. 3.3μL capsaicin | Wilcoxon test | two-tailed | ** | 0.0078 |

| | | | | | | |
|---|---|---|---|---|---|---|
| Fig.2C | Disgust Capsaicin | RM one-way ANOVA | 3.3μL vs. 33μL | Tukey's | * | 0.0137 |
| | | F (1.572, 11.00) = 30.66 | 3.3μL vs. 330μL | Tukey's | **** | <0.0001 |
| | | ****P<0.0001 | 33μL vs. 330μL | Tukey's | * | 0.0254 |
| | Disgust Quinine | RM one-way ANOVA | 100μL vs. 1000μL | Tukey's | ns | 0.4991 |
| | | F (1.322, 6.611) = 49.65 | 100μL vs. 10000μL | Tukey's | *** | 0.0002 |
| | | ***P=0.0002 | 1000μL vs. 10000μL | Tukey's | *** | 0.0005 |
| | Disgust Citric acid | RM one-way ANOVA | 10000μL vs. 100000μL | Tukey's | ns | 0.0975 |
| | | F (1.508, 7.538) = 86.24 | 10000μL vs. 1000000μL | Tukey's | *** | 0.0004 |
| | | ****P<0.0001 | 100000μL vs. 1000000μL | Tukey's | *** | 0.0004 |

| | | | | | | | |
|---|---|---|---|---|---|---|---|
| Fig.3C | Δdisgust vs. Δliking | 3.3μL vs. 33μL | Correlation | Two-tailed | ns | r=0.02287 | P=0.9571 |
| Fig.3D | | 10000μL vs. 100000μL | Correlation | Two-tailed | ns | r=-0.1962 | P=0.7095 |
| Fig.3E | | 33μL vs. 330μL | Correlation | Two-tailed | ns | r=0.5855 | P=0.1273 |
| Fig.3F | | 100000μL vs. 1000000μL | Correlation | Two-tailed | ns | r=0.3097 | P=0.5503 |

| | | | | | | | |
|---|---|---|---|---|---|---|---|
| Fig.4C | Tongue protrusion | Capsaicin | RM one-way ANOVA | 3.3μL vs. 33μL | Tukey's | ns | 0.9254 |
| | | | F (1.835, 12.84) = 7.153 | 3.3μL vs. 330μL | Tukey's | * | 0.0188 |
| | | | **P=0.0092 | 33μL vs. 330μL | Tukey's | * | 0.0284 |
| | | Quinine | RM one-way ANOVA | 100μL vs. 1000μL | Tukey's | ns | 0.335 |
| | | | F (1.142, 5.711) = 2.059 | 100μL vs. 10000μL | Tukey's | ns | 0.3813 |
| | | | P=0.2065 | 1000μL vs. 10000μL | Tukey's | ns | 0.6083 |
| | | Citric acid | RM one-way ANOVA | 10000μL vs. 100000μL | Tukey's | ns | 0.6257 |
| | | | F (1.757, 8.784) = 3.952 | 10000μL vs. 1000000μL | Tukey's | ns | 0.1655 |
| | | | P=0.0635 | 100000μL vs. 1000000μL | Tukey's | ns | 0.112 |
| Fig.4D | Gape | Capsaicin | RM one-way ANOVA | 3.3μL vs. 33μL | Tukey's | * | 0.0206 |
| | | | F (1.535, 10.74) = 4.784 | 3.3μL vs. 330μL | Tukey's | ns | 0.073 |
| | | | *P=0.0398 | 33μL vs. 330μL | Tukey's | ns | 0.9493 |
| | | Quinine | RM one-way ANOVA | 100μL vs. 1000μL | Tukey's | ns | 0.091 |
| | | | F (1.142, 5.710) = 2.853 | 100μL vs. 10000μL | Tukey's | ns | 0.2072 |
| | | | P=0.1443 | 1000μL vs. 10000μL | Tukey's | ns | 0.5279 |
| | | Citric acid | RM one-way ANOVA | 10000μL vs. 100000μL | Tukey's | ** | 0.0083 |
| | | | F (1.302, 6.511) = 24.34 | 10000μL vs. 1000000μL | Tukey's | ** | 0.0067 |
| | | | **P=0.0015 | 100000μL vs. 1000000μL | Tukey's | * | 0.0272 |
| | Chin rub | Capsaicin | RM one-way ANOVA | 3.3μL vs. 33μL | Tukey's | ns | 0.4526 |
| | | | F (1.937, 13.56) = 33.13 | 3.3μL vs. 330μL | Tukey's | *** | 0.0005 |
| | | | ****P<0.0001 | 33μL vs. 330μL | Tukey's | ** | 0.001 |
| | | Quinine | RM one-way ANOVA | 100μL vs. 1000μL | Tukey's | ns | 0.6083 |
| | | | F (1.937, 13.56) = 33.13 | 100μL vs. 10000μL | Tukey's | * | 0.0161 |
| | | | ****P<0.0001 | 1000μL vs. 10000μL | Tukey's | * | 0.0184 |
| | | Citric acid | RM one-way ANOVA | 10000μL vs. 100000μL | Tukey's | Not analyzed (NA) | |
| | | | F (1.000, 5.000) = 49.47 | 10000μL vs. 1000000μL | Tukey's | ** | 0.0021 |
| | | | **P=0.0009 | 100000μL vs. 1000000μL | Tukey's | ** | 0.0021 |
| | Head shake | Capsaicin | RM one-way ANOVA | 3.3μL vs. 33μL | Tukey's | ns | >0.9999 |
| | | | F (1.471, 10.29) = 0.4667 | 3.3μL vs. 330μL | Tukey's | ns | 0.6 |
| | | | P=0.5825 | 33μL vs. 330μL | Tukey's | ns | 0.6 |
| | | Quinine | RM one-way ANOVA | 100μL vs. 1000μL | Tukey's | ns | >0.9999 |
| | | | F (1.220, 6.098) = 4.000 | 100μL vs. 10000μL | Tukey's | ns | 0.0552 |
| | | | P=0.0881 | 1000μL vs. 10000μL | Tukey's | ns | 0.0552 |
| | | Citric acid | RM one-way ANOVA | 10000μL vs. 100000μL | Tukey's | ns | 0.2779 |
| | | | F (1.000, 5.000) = 3.049 | 10000μL vs. 1000000μL | Tukey's | ns | 0.2779 |
| | | | P=0.1412 | 100000μL vs. 1000000μL | Tukey's | NA | |
| | Face wash | Capsaicin | RM one-way ANOVA | 3.3μL vs. 33μL | Tukey's | * | 0.0363 |
| | | | F (1.712, 11.99) = 10.69 | 3.3μL vs. 330μL | Tukey's | ** | 0.0039 |
| | | | **P=0.0028 | 33μL vs. 330μL | Tukey's | ns | 0.3269 |
| | | Quinine | RM one-way ANOVA | 100μL vs. 1000μL | Tukey's | ns | >0.9999 |
| | | | F (1.724, 8.619) = 13.62 | 100μL vs. 10000μL | Tukey's | ** | 0.009 |
| | | | **P=0.0026 | 1000μL vs. 10000μL | Tukey's | ** | 0.009 |
| | | Citric acid | RM one-way ANOVA | 10000μL vs. 100000μL | Tukey's | ns | 0.6083 |
| | | | F (1.012, 5.060) = 212.6 | 10000μL vs. 1000000μL | Tukey's | **** | <0.0001 |
| | | | ****P<0.0001 | 100000μL vs. 1000000μL | Tukey's | **** | <0.0001 |
| | Forelimb flail | Capsaicin | RM one-way ANOVA | 3.3μL vs. 33μL | Tukey's | ** | 0.0085 |
| | | | F (1.610, 11.27) = 17.47 | 3.3μL vs. 330μL | Tukey's | ** | 0.0015 |
| | | | ***P=0.0006 | 33μL vs. 330μL | Tukey's | ns | 0.1401 |
| | | Quinine | RM one-way ANOVA | 100μL vs. 1000μL | Tukey's | ns | 0.4482 |
| | | | F (1.331, 6.654) = 40.88 | 100μL vs. 10000μL | Tukey's | *** | 0.0005 |
| | | | ***P=0.0003 | 1000μL vs. 10000μL | Tukey's | *** | 0.0005 |

| | | | | | | |
|---|---|---|---|---|---|---|
| | | Citric acid | RM one-way ANOVA<br>F (1.350, 6.750) = 36.67<br>***P=0.0004 | 10000μL vs. 100000μL | Tukey's | ns | 0.1897 |
| | | | | 10000μL vs. 1000000μL | Tukey's | ** | 0.0034 |
| | | | | 100000μL vs. 1000000μL | Tukey's | ** | 0.0026 |

| | | | | | | |
|---|---|---|---|---|---|---|
| Supplemental fig.1A | Neutral<br>solution type | RM one-way ANOVA<br>F (1.434, 8.601) = 2.563<br>P=0.1410 | Water vs. 0.05%EtOH | Tukey's | ns | 0.991 |
| | | | Water vs. 3.3 | Tukey's | ns | 0.059 |
| | | | 0.05%EtOH vs. 3.3 | Tukey's | ns | 0.2126 |
| Supplemental fig.1B | Disgust<br>solution type | RM one-way ANOVA<br>F (1.219, 7.316) = 0.2898<br>P=0.6510 | Water vs. 0.05%EtOH | Tukey's | ns | 0.8827 |
| | | | Water vs. 3.3 | Tukey's | ns | 0.8346 |
| | | | 0.05%EtOH vs. 3.3 | Tukey's | ns | 0.8843 |
| Supplemental fig.1C | Neutral<br>Hot water | RM one-way ANOVA<br>F (2.515, 15.09) = 7.617<br>**P=0.0034 | Water vs. 50°C | Tukey's | * | 0.0364 |
| | | | Water vs. 70°C | Tukey's | ** | 0.0045 |
| | | | Water vs. 90°C | Tukey's | ns | 0.4078 |
| | | | 50°C vs. 70°C | Tukey's | ns | 0.7982 |
| | | | 50°C vs. 90°C | Tukey's | ns | 0.4082 |
| | | | 70°C vs. 90°C | Tukey's | ns | 0.1963 |
| Supplemental fig.1D | Disgust<br>Hot water | RM one-way ANOVA<br>F (1.489, 8.931) = 2.350<br>P=0.1568 | Water vs. 50°C | Tukey's | ns | 0.9713 |
| | | | Water vs. 70°C | Tukey's | ns | 0.5283 |
| | | | Water vs. 90°C | Tukey's | ns | 0.4054 |
| | | | 50°C vs. 70°C | Tukey's | ns | 0.379 |
| | | | 50°C vs. 90°C | Tukey's | ns | 0.2065 |
| | | | 70°C vs. 90°C | Tukey's | ns | 0.9926 |
| Supplemental fig.1E | Neutral | CPZ vs. DMSO | Wilcoxon test | two-tailed | ns | 0.1094 |
| Supplemental fig.1F | Disgust | CPZ vs. DMSO | Wilcoxon test | two-tailed | ns | >0.9999 |

| | | | | | | |
|---|---|---|---|---|---|---|
| Supplemental fig.4B | Neutral<br>Capsaicin | RM one-way ANOVA<br>F (1.699, 11.89) = 45.32<br>****P<0.0001 | 3.3μL vs. 33μL | Tukey's | ns | 0.4132 |
| | | | 3.3μL vs. 330μL | Tukey's | *** | 0.0001 |
| | | | 33μL vs. 330μL | Tukey's | **** | <0.0001 |
| | Neutral<br>Quinine | RM one-way ANOVA<br>F (1.449, 7.247) = 15.05<br>**P=0.0038 | 100μL vs. 1000μL | Tukey's | ns | 0.7668 |
| | | | 100μL vs. 10000μL | Tukey's | ** | 0.0017 |
| | | | 1000μL vs. 10000μL | Tukey's | * | 0.0168 |
| | Neutral<br>Citric acid | RM one-way ANOVA<br>F (1.362, 6.809) = 2.036<br>P=0.2019 | 10000μL vs. 100000μL | Tukey's | ns | 0.445 |
| | | | 10000μL vs. 1000000μL | Tukey's | ns | 0.8556 |
| | | | 100000μL vs. 1000000μL | Tukey's | ns | 0.0617 |
| Supplemental fig.4D | Mouth movements<br>Capsaicin | RM one-way ANOVA<br>F (1.699, 11.89) = 45.32<br>****P<0.0001 | 3.3μL vs. 33μL | Tukey's | ns | 0.4132 |
| | | | 3.3μL vs. 330μL | Tukey's | *** | 0.0001 |
| | | | 33μL vs. 330μL | Tukey's | **** | <0.0001 |
| | Mouth movements<br>Quinine | RM one-way ANOVA<br>F (1.449, 7.247) = 15.05<br>**P=0.0038 | 100μL vs. 1000μL | Tukey's | ns | 0.7668 |
| | | | 100μL vs. 10000μL | Tukey's | ** | 0.0017 |
| | | | 1000μL vs. 10000μL | Tukey's | * | 0.0168 |
| | Mouth movements<br>Citric acid | RM one-way ANOVA<br>F (1.362, 6.809) = 2.036<br>P=0.2019 | 10000μL vs. 100000μL | Tukey's | ns | 0.445 |
| | | | 10000μL vs. 1000000μL | Tukey's | ns | 0.8556 |
| | | | 100000μL vs. 1000000μL | Tukey's | ns | 0.0617 |

| | | | | | | |
|---|---|---|---|---|---|---|
| Supplemental fig.5 | Gape<br>Ratio<br>Capsaicin | RM one-way ANOVA<br>F (1.126, 7.880) = 2.838<br>P=0.1297 | 3.3μL vs. 33μL | Tukey's | ns | 0.6508 |
| | | | 3.3μL vs. 330μL | Tukey's | ns | 0.0949 |
| | | | 33μL vs. 330μL | Tukey's | ns | 0.1387 |
| | Gape<br>Ratio<br>Quinine | RM one-way ANOVA<br>F (1.067, 5.334) = 0.8507<br>P=0.4042 | 100μL vs. 1000μL | Tukey's | ns | 0.8301 |
| | | | 100μL vs. 10000μL | Tukey's | ns | 0.5948 |
| | | | 1000μL vs. 10000μL | Tukey's | ns | 0.1687 |
| | Gape<br>Ratio<br>Citric acid | RM one-way ANOVA<br>F (1.187, 5.936) = 5.113<br>P=0.0616 | 10000μL vs. 100000μL | Tukey's | ns | 0.9651 |
| | | | 10000μL vs. 1000000μL | Tukey's | ns | 0.1721 |
| | | | 100000μL vs. 1000000μL | Tukey's | ns | 0.1094 |
| | Chin rub<br>Ratio<br>Capsaicin | RM one-way ANOVA<br>F (1.290, 9.031) = 3.003<br>P=0.1120 | 3.3μL vs. 33μL | Tukey's | ns | 0.8076 |
| | | | 3.3μL vs. 330μL | Tukey's | ns | 0.3432 |
| | | | 33μL vs. 330μL | Tukey's | ** | 0.0077 |
| | Chin rub<br>Ratio<br>Quinine | RM one-way ANOVA<br>F (1.290, 9.031) = 3.003<br>P=0.1120 | 100μL vs. 1000μL | Tukey's | ns | 0.6083 |
| | | | 100μL vs. 10000μL | Tukey's | * | 0.0164 |
| | | | 1000μL vs. 10000μL | Tukey's | ns | 0.0518 |
| | Chin rub<br>Ratio<br>Citric acid | RM one-way ANOVA<br>F (1.000, 5.000) = 22.06<br>**P=0.0054 | 10000μL vs. 100000μL | Tukey's | NA | |
| | | | 10000μL vs. 1000000μL | Tukey's | * | 0.0123 |
| | | | 100000μL vs. 1000000μL | Tukey's | * | 0.0123 |
| | Head shake<br>Ratio<br>Capsaicin | RM one-way ANOVA<br>F (1.063, 7.442) = 0.7703<br>P=0.4157 | 3.3μL vs. 33μL | Tukey's | ns | 0.7467 |
| | | | 3.3μL vs. 330μL | Tukey's | ns | 0.6 |
| | | | 33μL vs. 330μL | Tukey's | ns | 0.6 |
| | Head shake<br>Ratio<br>Quinine | RM one-way ANOVA<br>F (1.230, 6.151) = 1.304<br>P=0.3098 | 100μL vs. 1000μL | Tukey's | ns | 0.9933 |
| | | | 100μL vs. 10000μL | Tukey's | ns | 0.2391 |
| | | | 1000μL vs. 10000μL | Tukey's | ns | 0.2914 |
| | Head shake<br>Ratio<br>Citric acid | RM one-way ANOVA<br>F (1.000, 5.000) = 3.195<br>P=0.1339 | 10000μL vs. 100000μL | Tukey's | ns | 0.265 |
| | | | 10000μL vs. 1000000μL | Tukey's | ns | 0.265 |
| | | | 100000μL vs. 1000000μL | Tukey's | NA | |
| | Face wash<br>Ratio<br>Capsaicin | RM one-way ANOVA<br>F (1.370, 9.587) = 0.2170<br>P=0.7264 | 3.3μL vs. 33μL | Tukey's | ns | 0.8468 |
| | | | 3.3μL vs. 330μL | Tukey's | ns | 0.9126 |
| | | | 33μL vs. 330μL | Tukey's | ns | 0.9653 |
| | Face wash<br>Ratio<br>Quinine | RM one-way ANOVA<br>F (1.370, 6.851) = 0.8199<br>P=0.4349 | 100μL vs. 1000μL | Tukey's | ns | 0.5432 |
| | | | 100μL vs. 10000μL | Tukey's | ns | 0.7264 |
| | | | 1000μL vs. 10000μL | Tukey's | ns | 0.9017 |
| | Face wash<br>Ratio<br>Citric acid | RM one-way ANOVA<br>F (1.146, 5.730) = 4.894<br>P=0.0685 | 10000μL vs. 100000μL | Tukey's | ns | 0.4716 |
| | | | 10000μL vs. 1000000μL | Tukey's | ns | 0.4301 |
| | | | 100000μL vs. 1000000μL | Tukey's | *** | 0.0009 |
| | Forelimb flail<br>Ratio<br>Capsaicin | RM one-way ANOVA<br>F (1.115, 7.802) = 6.331<br>*P=0.0344 | 3.3μL vs. 33μL | Tukey's | ns | 0.0514 |
| | | | 3.3μL vs. 330μL | Tukey's | ns | 0.1651 |
| | | | 33μL vs. 330μL | Tukey's | ns | 0.0825 |
| | Forelimb flail<br>Ratio<br>Quinine | RM one-way ANOVA<br>F (1.096, 5.482) = 1.679<br>P=0.2509 | 100μL vs. 1000μL | Tukey's | ns | 0.5536 |
| | | | 100μL vs. 10000μL | Tukey's | ns | 0.3982 |
| | | | 1000μL vs. 10000μL | Tukey's | ns | 0.3359 |
| | Forelimb flail<br>Ratio<br>Citric acid | RM one-way ANOVA<br>F (1.620, 8.098) = 3.030<br>P=0.1097 | 10000μL vs. 100000μL | Tukey's | ns | 0.268 |
| | | | 10000μL vs. 1000000μL | Tukey's | ns | 0.1984 |
| | | | 100000μL vs. 1000000μL | Tukey's | ns | 0.558 |

**Supplemental figure 1**

**Mild higher temperature (50-70˚C) increases neutral reactions**

**(A,B)** Scores for neutral reactions (**A**) and disgust reactions (**B**) during intraoral infusion of water, vehicle solution (0.05% ethanol), and 3.3 µM capsaicin solution (*n*=7 mice from two independent experiments). The black circles represent individual measurements from single injections.

**(C,D)** Scores for neutral reactions (**C**) and disgust reactions (**D**) during intraoral infusion of water at room temperature, 50 °C, 70 °C, and 90 °C (*n*=7 mice from two independent experiments). The black circles represent individual measurements from single injections.

**(E,F)** Scores for neutral reactions (**E**) and disgust reactions (**F**) during intraoral infusion of water in mice injected with capsazepine or vehicle (*n*=7 mice from two independent experiments). The black circles represent individual measurements from single injections.

**Supplemental figure 2**

**Representative time-lapse sequences of disgust reactions during intraoral infusion of quinine**

The numbers at the bottom of each panel indicate the elapsed time (milliseconds). G, gape; CR, chin rub; HS, head shake; FW, face wash; FF, forelimb flail. Scale bar, 5 mm.

**Supplemental figure 3**

**Corresponding to Fig. 4C, D**

The scores for tongue protrusion (**A**), gape, chin rub, head shake, face wash, and forelimb flail (**B**) during intraoral infusion of three different concentrations of capsaicin (left panel; *n* = 8 mice from six independent experiments), quinine (middle panel; *n* = 6 mice from four independent experiments), and citric acid (right panel; *n* = 6 mice from four independent experiments).

Error bars are SEM.

**Supplemental figure 4**

**Capsaicin-induced neutral reactions decrease as capsaicin concentration increase**

**(A)** Relationship between the score of neutral reactions during intraoral infusion of capsaicin (red circles; *n* = 8 mice from six independent experiments), quinine (green rectangles; $n$ = 6 mice from four independent experiments), and citric acid (blue triangles; $n$ = 6 mice from four independent experiments), and their concentrations.

**(B)** The score for neutral reactions during intraoral infusion of three different concentrations of capsaicin (left panel; $n$ = 8 mice from six independent experiments), quinine (middle panel; $n$ = 6 mice from four independent experiments), and citric acid (right panel; $n$ = 6 mice from four independent experiments). The black circles represent individual measurements from single injections.

**(C)** Relationship between the score of mouth movements during intraoral infusion of capsaicin (red circles; $n$ = 8 mice from six independent experiments), quinine (green rectangles; $n$ = 6 mice from four independent experiments), and citric acid (blue triangles; $n$ = 6 mice from four independent experiments), and their concentrations.

**(D)** Score for mouth movements during intraoral infusion of three different concentrations of capsaicin ($n$ = 8 mice from six independent experiments), quinine ($n$ = 6 mice from four independent experiments), and citric acid ($n$ = 6 mice from four independent experiments). The black circles represent individual measurements from single injections.

Error bars are SEM. *$P$ < .05, **$P$ < .01, ***$P$ < four001, ****$P$ < .0001, Tukey's test.

**Supplemental figure 5**

**Corresponding to Fig. 6A-E**

The ratio of gape, chin rub, head shake, face wash, and forelimb flail among all disgust reactions induced by capsaicin ($n$ = 8 mice from six independent experiments), quinine ($n$ = 6 mice from four independent experiments), and citric acid ($n$ = 6 mice from four independent experiments). The ratio of chin rub significantly increased at high concentrations of capsaicin, quinine, and citric acid. The black circles represent individual measurements from single injections.

*$P$ < .05, **$P$ < .01, Tukey's test.
